# A Novel *C. elegans* Model for Tau Spreading Reveals Genes Critical for Endolysosomal Integrity and Seeded Tau Aggregation

**DOI:** 10.1101/2024.11.13.619586

**Authors:** Carl Alexander Sandhof, Nicole Martin, Jessica Tittelmeier, Annabelle Schlueter, Martina Pezzali, David C. Schoendorf, Timo Lange, Peter Reinhardt, Janina S. Ried, Siwen Liang, Gamze Uzunoglu, Laura Gasparini, Thomas R. Jahn, Dagmar E. Ehrnhoefer, Carmen Nussbaum-Krammer

**Affiliations:** Center for Molecular Biology of Heidelberg University (ZMBH) and German Cancer Research Center (DKFZ), DKFZ-ZMBH Alliance, Im Neuenheimer Feld 282, 69120 Heidelberg, Germany; Chemical Neurobiology Laboratory, Center for Genomic Medicine, Departments of Neurology and Psychiatry, Massachusetts General Hospital and Harvard Medical School, Boston, MA 02114, USA; Chair of Neuroanatomy, Institute of Anatomy, Faculty of Medicine, LMU Munich, Pettenkoferstrasse 11, 80336 Munich, Germany; AbbVie Deutschland GmbH & Co. KG, Neuroscience Discovery, Knollstrasse 50, 67061 Ludwigshafen am Rhein, Germany; AbbVie Deutschland GmbH & Co. KG, Genomics Research Center, Knollstrasse 50, 67061 Ludwigshafen am Rhein, Germany

**Keywords:** Alzheimer’s disease, tauopathies, prion-like spreading, neurodegeneration, endolysosomal system, lysosomal membrane permeabilization, induced pluripotent stem cells

## Abstract

The spreading of Tau pathology is closely associated with the progression of neurodegeneration and cognitive decline in Alzheimer’s disease and other tauopathies. A key event in this process is the rupture of endolysosomal vesicles following the intercellular transfer of Tau aggregates, releasing the transferred Tau species into the cytosol where they can promote the aggregation of endogenous Tau. However, understanding of the cellular pathways involved in this process remains limited.

In this study, we investigated cellular pathways that prevent endolysosomal vesicle rupture. We established a new *C. elegans* model of Tau spreading by introducing an mCherry-labelled, disease-associated aggregation-prone fragment of human Tau (F3^ΔK281^::mCh) into the six touch receptor neurons. F3^ΔK281^::mCh transgenic animals exhibited significant neurotoxicity and mechanosensory deficits due to the accumulation of this Tau fragment. In addition, its intercellular transmission compromised the endolysosomal system in receiving hypodermal cells. Using this model, we conducted an unbiased genome-wide RNAi screen and identified 59 genes critical for maintaining endolysosomal integrity. GO-term analysis revealed an enrichment of genes related to the ESCRT complex, the ubiquitin-proteasome system, mRNA splicing, and fatty acid metabolism. Silencing of selected conserved genes exacerbated seeded Tau aggregation in a human induced pluripotent stem cell (hiPSC)-derived cortical neuron model and triggered endolysosomal rupture in HEK293T cells, confirming the crucial role of endolysosomal damage in seeded Tau aggregation.

Overall, this study discovered novel cellular pathways that safeguard endolysosomal integrity. These findings may guide the development of therapeutics that improve endolysosomal integrity to halt the progression of Tau pathology.

## Introduction

Age-related neurodegenerative diseases, such as Alzheimer’s disease (AD) and Parkinson’s disease (PD), are characterized by the gradual accumulation of amyloid deposits that contain fibrillar aggregates of disease specific proteins. Aggregation of the microtubule-associated protein Tau (MAPT/Tau) is associated with AD and other tauopathies, whereas α-synuclein (SNCA/α-Syn) containing inclusions are associated with PD and other synucleinopathies [1].

These pathological inclusions spread from an initial site of origin to functionally interconnected brain regions, which correlates with the progression of neurodegeneration and cognitive decline [2–4]. The molecular basis of this progressive spreading behavior lies in the unique structure of amyloid fibrils, which are composed of highly ordered β-sheet-rich conformations of the disease protein. Amyloid fibrils can self-propagate by a process known as seeded aggregation, in which the structure of the amyloid fibril acts as a template that drives the conversion and incorporation of the soluble native protein [3,5]. In addition, small fibrillar fragments (seeds) can invade and initiate the aggregation of native proteins in neighboring cells, accounting for the continuing spreading of pathology [3,5]. Although neuronal death is central to these diseases, the pathology is not limited to neurons but extends to non-neuronal cell types such as glial cells, suggesting a broader cellular involvement in the pathogenesis of these disorders [1].

The intercellular propagation of protein misfolding involves several key steps. First, disease-associated proteins are released from one cell (donor cell) and subsequently taken up by neighboring or synaptically connected cells (receiving cells). Internalization of Tau and α-Syn is often mediated by endocytosis [3,6]. Once in the recipient cell, these misfolded proteins seed the aggregation of native proteins [10,11]. However, for seeded aggregation to occur, pathological proteins must first escape from endolysosomal vesicles and access the cytosol, where they can interact with native proteins and induce their aggregation. This escape from the endolysosomal vesicles appears to be the rate-limiting step for the aggregation of native proteins in the receiving cells [10,11]. Thus, successful intercellular spreading of Tau pathology depends on the ability of the transmitted protein species to evade degradation and exit endolysosomal vesicles. Pathological Tau and α-Syn are able to damage endo-membranes, facilitating their escape from endolysosomal vesicles [7–9]. Unraveling the pathways involved in endolysosomal escape is crucial to elucidating the mechanisms driving the spreading of Tau pathology. Moreover, cellular factors that protect endolysosomal integrity may not only be relevant to tauopathies, but also to other age-related diseases and aging in general.

The nematode *Caenorhabditis elegans (C. elegans)* has been instrumental in discovering genetic modifiers of neurodegenerative diseases, including AD and tauopathies, and to study the intercellular spreading of proteins associated with neurodegenerative diseases [12,13]. We have previously shown that expression as well as the chronic intercellular transfer of misfolded α-Syn causes endolysosomal damage in donor and receiving cells [14], consistent with observations in human neurons [7,9]. We also found that reducing DAF-2 insulin-like signaling, which is well known for its capacity to extend lifespan, effectively prevented endolysosomal damage [14]. This indicates that slowing aging processes may protect endolysosomal integrity and thereby reduce α-Syn propagation and toxicity. Nevertheless, comprehensive exploration of additional cellular pathways that contribute to the maintenance of endolysosomal integrity awaits further investigation.

In this study, we generated and characterized a new complementary *C. elegans* model for Tau spreading. We expressed a highly amyloidogenic fragment of human Tau (F3) harboring a disease-associated mutation (ΔK281) [15,16] fused to mCherry (F3^ΔK281^::mCh) in the six touch receptor neurons of *C. elegans*. F3^ΔK281^::mCh transgenic animals displayed prominent neurotoxicity and mechanosensory deficits. Moreover, intercellular transmission of Tau compromised the receiving hypodermal endolysosomal system. Using this strain, we performed an unbiased genome-wide RNAi screen for modifiers of endolysosomal rupture, identifying 59 genes critical for maintaining endolysosomal integrity. Gene Ontology (GO)-term analysis showed an enrichment of genes associated with the ESCRT complex, the ubiquitin-proteasome system, mRNA splicing, and fatty acid metabolism. To validate our *in vivo* findings in *C. elegans*, we examined the effect of the identified genes in human cells. As expected, knockdown of selected conserved genes induced endolysosomal rupture in HEK293T cells and exacerbated seeded Tau aggregation in a human induced pluripotent stem cell (hiPSC)-derived cortical neuron model.

Overall, this study shed light on the relationship between endolysosomal integrity and seeded Tau aggregation and revealed cellular pathways involved. These results may provide additional insights regarding preserving endolysosomal integrity and ameliorating the spreading of Tau pathology in AD and other tauopathies.

## Results

### Expression of F3^ΔK281^::mCh in touch receptor neurons causes neurotoxicity

*C. elegans* is an excellent model organism for the discovery and characterization of cellular pathways and genetic modifiers implicated in the intercellular propagation of disease-related proteins [17]. We have previously developed two *C. elegans* models to study basic mechanisms of α-Syn spreading [14]. Tracking RFP-labeled α-Syn, selectively expressed in either body wall muscle (BWM) or dopaminergic (DA) neurons revealed its transmission to the adjacent hypodermis [14]. The use of these model systems allowed the identification and characterization of genes that modulate this step in α-Syn::RFP spreading, contributing to a better understanding of the pathways involved in intercellular transmission of α-Syn [14,18].

Since Tau also shows spreading behavior [2–4], we aimed to generate a corresponding *C. elegans* model for Tau spreading. The sole nematode homolog of Tau, called protein with Tau-like repeats (PTL-1), is predominantly expressed in touch receptor neurons [19,20]. Hence, we expressed Tau in these six neurons by using the promoter of the *mec-4* gene, encoding an ion channel protein that is exclusively active in the touch receptor neurons [21] (Figure 1A). The expression plasmid contains an mCh-labelled, aggregation prone fragment (F3) of human Tau harboring a mutation (ΔK281, previously named ΔK280) associated with familial forms of frontotemporal dementia [16,22] (Figure 1B), a primary tauopathy. The F3 fragment was discovered in a cell culture model as a product of sequential proteolytic cleavage of the repeat domain of Tau, which forms the core of Tau amyloids [15]. Pan-neuronal expression of this highly aggregation-prone fragment resulted in severe neurotoxicity in a previously established *C. elegans* model [16].

**Figure 1.**
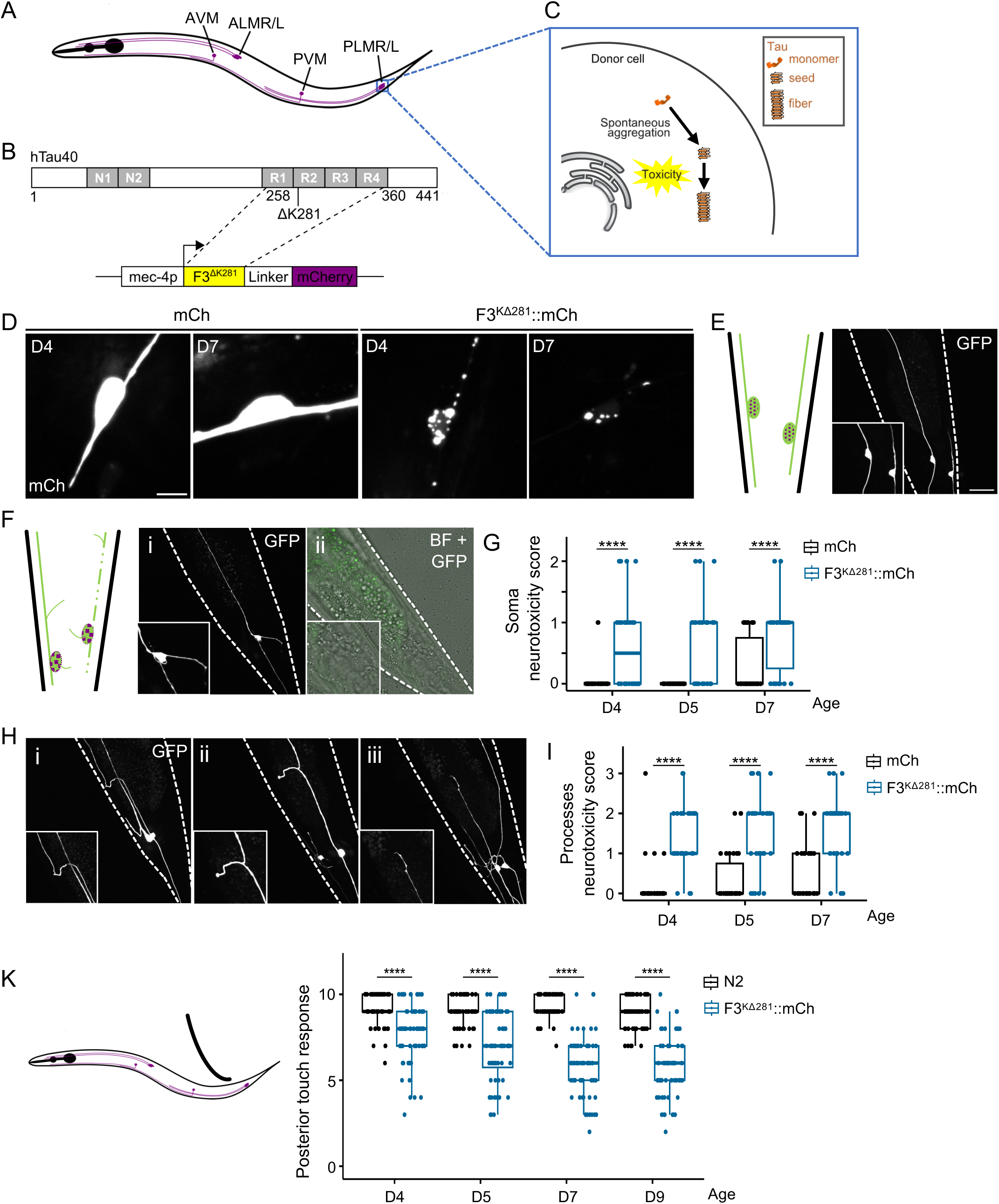
Expression of an aggregation prone Tau fragment in touch receptor neurons of *C. elegans* is neurotoxic. (A) Schematic illustration of median plane representation of the six touch receptor neurons and their processes in an adult *C. elegans* hermaphrodite. AVM: anterior ventral microtubule cell; ALMR/L: anterior lateral microtubule cell right/left; PVM: posterior ventral microtubule cell; PLMR/L: posterior lateral microtubule cell right/left. (B) Schematic representation of the full-length human Tau 2N4R splice variant with the frontotemporal dementia-associated K281 deletion and the expression construct generated to express the F3^ΔK281^ fragment fused to a C-terminal mCherry (mCh) tag under the touch receptor neuron specific *mec-4* promotor. (C) Schematic illustration of monomeric Tau spontaneously aggregating into seeds and fibers. The toxicity of F3^ΔK281^::mCh was assessed in the PLML/R neurons (donor cell). (D) Collapsed confocal z-stacks (561 nm channel) of the tail region of animals expressing mCh control or the F3^ΔK281::^mCh Tau fragment alongside a GFP marker. Animals were analyzed at day 4 (D4) and day 7 (D7) of age. Foci indicate F3^ΔK281^::mCh aggregation. Scale bar = 5 µm. (E, F, H) Collapsed confocal z-stacks (488 nm channel) of 7-days-old animals expressing F3^ΔK281^::mCh and GFP in touch receptor neurons. Scale bar = 20 μm. (E) Healthy PLM neurons exhibit intact soma and straight neuronal processes. (F) PLM neuron toxicity phenotypes observed in the soma include (i) abnormal soma outgrowth and (ii) neurodegeneration, indicated by missing PLM soma. (H) Toxicity phenotypes in PLM processes were classified as (i) wavy, (ii) branching and/or (iii) broken or prematurely ending processes. (G, I, K) Quantitative analysis of neurotoxicity phenotypes. (G) PLM neurons expressing F3^ΔK281::^mCh exhibit significantly higher soma neurotoxicity compared to the mCh control. (I) PLM processes also show significantly more toxicity phenotypes upon F3^ΔK281^::mCh expression compared to the mCh control. (K) Left: Schematic showing the region of gentle touch stimulation used to assess posterior touch responses. Right: Quantification of posterior touch responses (out of 10 stimuli) in wild-type N2 control and F3^ΔK281^::mCh expressing animals. Each point represents the response of a single animal. In panels (G, I, K) data are presented as boxplot, showing the mean, upper and lower quartiles, and the minimum and maximum values. Points outside the min-max range represent outliers. N = 3 with 10-20 animals per strain, timepoint and replicate. Statistical analysis was conducted using Two-Way mixed-model ANOVA on rank-transformed data, with pairwise comparisons of estimated marginal means with Bonferroni correction for multiple comparisons. **** = p < 0.0001.

We first assessed the aggregation and toxicity of F3^ΔK281^::mCh (Figure 1C). Consistent with the high aggregation propensity of the F3^ΔK281^ fragment, its expression in touch receptor neurons resulted in the formation of numerous intracellular foci as opposed to the diffuse distribution of the mCh tag-only control (Figure 1D). To examine the toxicity of the Tau F3^ΔK281^ fragment in our model, we monitored the integrity of the PLM neuronal soma and processes from day four to day seven of life (Figure 1E-I). To ensure robust monitoring of neuronal integrity, we co-expressed a GFP marker in the six touch receptor neurons in addition to F3^ΔK281^::mCh or mCh. Healthy PLM neurons send a straight, unbranched process to the anterior and a shorter process posteriorly (Figure 1E). Upon expression of F3^ΔK281^::mCh, PLM neurons exhibited additional, abnormal soma outgrowths (Figure 1Fi) or neurodegeneration, as indicated by a missing soma (Figure 1Fii). At all ages assessed, these abnormal soma phenotypes were significantly more frequent in F3^ΔK281^::mCh expressing animals compared to the mCh control (Figure 1G). In addition, F3^ΔK281^::mCh expression caused a wavy appearance (Figure 1Hi), branching events (Figure 1Hii) and/or breaks (Figure 1Hiii) in the PLM processes to a greater extent than the mCh control (Figure 1I).

As a consequence of this neurotoxicity, mechano-sensation of F3^ΔK281^::mCh expressing animals was impaired, leading to an age-dependent decline in the response to a posterior touch stimulus (Figure 1K).

The F3^ΔK281^ fragment acts as a nucleation agent and promotes aggregation of full-length wild-type Tau [15]. Therefore, we asked whether it could also affect the Tau homolog PTL-1 in *C. elegans* touch neurons. While we did not observe co-aggregation by microscopy, expression of F3^ΔK281^::mCh disrupted the association of PTL-1 with microtubules and led to an overall decrease in PTL-1 levels (Figure S1A, B). This was not due to a global negative effect on cellular protein levels as the fluorescence intensity of GFP did not significantly differ between F3^ΔK281^::mCh and mCh expressing animals (Figure S1C, D).

Taken together, expression of F3^ΔK281^::mCh causes neurotoxicity in *C. elegans.* Since it also disrupts PTL-1 localization and levels, the observed effects may result not only from a toxic gain of function due to the accumulation of F3^ΔK281^::mCh, but also from the loss of physiological function of the endogenous Tau homolog PTL-1 [23].

### F3^ΔK281^::mCh is transmitted to the hypodermis and damages the hypodermal endolysosomal system

Prion-like protein aggregates not only amplify within a cell but also disseminate to neighboring cells and tissues (Figure 2A). We have previously shown that α-Syn is transmitted from different tissues to the adjacent hypodermis in *C. elegans* in an age-dependent manner [14,18]. Since Tau also disseminates to neighboring cells in tauopathies, we examined whether F3^ΔK281^::mCh is also transferred to the hypodermis in this *C. elegans* Tau model. The projections of the touch receptor neurons are embedded within the hypodermis, which likely facilitates the transmission of F3^ΔK281^::mCh (Figure 2B). Indeed, fluorescent signals were detected as early as day one, the L1 larval stage (Figure 2C, D). The amount of transferred F3^ΔK281^::mCh increased with time and was significantly higher compared to the mCh control at all examined ages (Figure 2C, D).

**Figure 2.**
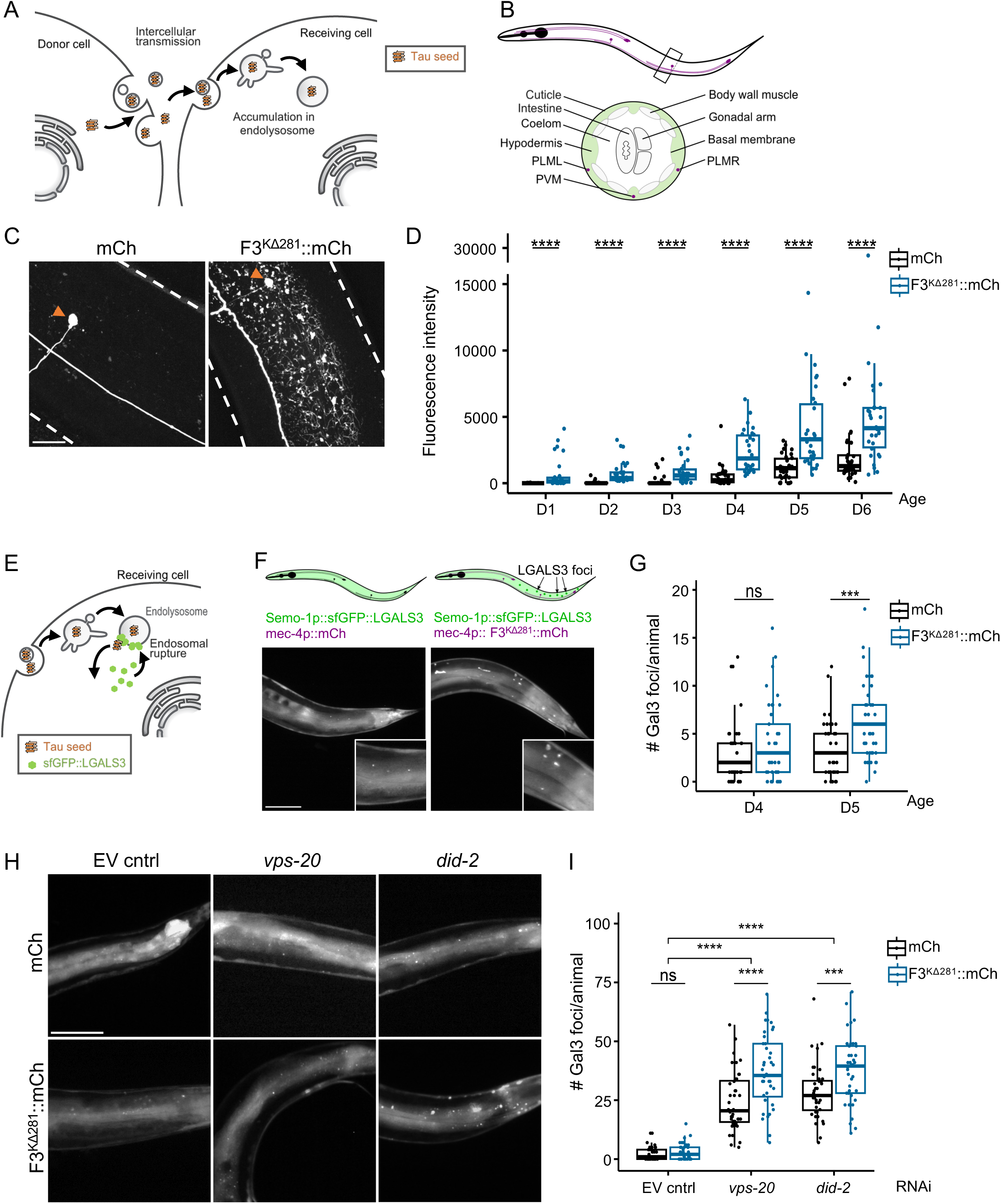
Transmission of the F3 Tau fragment from touch receptor neurons to the hypodermis affects the endolysosomal system. (A) Schematic illustration of the intercellular transmission of F3^ΔK281^::mCh from touch receptor neurons (donor cell) to hypodermal cells (receiving cell). (B) Top: Schematic median plane representation of the six F3^ΔK281^::mCh expressing touch receptor neurons and their processes in an adult *C. elegans* hermaphrodite. Bottom: Cross sectionals view of the area marked by a rectangle indicating the spatial relationship of the hypodermal tissue (green) and the processes of the posterior set of touch receptor neurons (purple). While the hypodermis is demarcated from most other tissues by a basal membrane the touch receptor neuron processes are directly embedded in it. (C) Collapsed confocal z-stack of 6-days-old nematodes expressing mCh or F3^ΔK281^::mCh in the touch receptor neurons. Worm boundaries are outlined by dashed lines with orange arrowheads indicating PVM soma. Scale bar = 20 µm. (D) Quantification of normalized fluorescence intensity of mCh and F3^ΔK281^::mCh in the hypodermis at indicated ages. (E) Schematic illustration of endolysosomal damage detection using the galectin-3 reporter (sfGFP::LGALS3) in receiving hypodermal cells. (F) Top: Schematics represent the sfGFP::LGALS3 signal distribution corresponding to the representative fluorescence widefield images below. Bottom: Representative widefield fluorescence images of 5-days-old animals expressing mCh control or F3^ΔK281^::mCh in touch receptor neurons and sfGFP::LGALS3 in the hypodermis. In control animals the sfGFP::LGALS3 (Gal3) signal in the hypodermis remains mostly diffuse (left), while expression of F3^ΔK281^::mCh in touch receptor neurons leads to hypodermal Gal3 foci formation indicating endolysosomal membrane rupture (right). Scale bar = 100 µm. (G) Quantification of hypodermal Gal3 foci in animals co-expressing mCh or F3^ΔK281^::mCh in touch receptor neurons. (H) Representative fluorescence widefield images of hypodermal Gal3 foci in the newly generated sfGFP::LGALS3 strain co-expressing mCh or F3^ΔK281^::mCh in touch receptor neurons, grown on empty vector (EV) control or indicated RNAi plates at day 5 of age. Scale bar = 100 µm. (I) Quantification of hypodermal Gal3 foci in the newly generated sfGFP::LGALS3 strain co-expressing mCh or F3^ΔK281^::mCh in touch receptor neurons, grown on EV control or indicated RNAi plates at day 5 of age. Co-expression of F3^ΔK281^::mCh alone does not significantly induce endolysosomal rupture but enhanced lysosomal rupture with RNAi knockdown of the ESCRT-III complex components *vps-20* or *did-2*. (D, G, I) Data are presented as boxplot, showing the mean, upper and lower quartiles, and the minimum and maximum values. Points outside the min-max range represent outliers. N = 3 with 10-15 animals per strain, timepoint and replicate. Each point represents one animal. Statistical analysis was conducted using Two-Way mixed-model ANOVA on rank-transformed data, with pairwise comparisons of estimated marginal means with Bonferroni correction for multiple comparisons. ns = not significant, *** = p < 0.001, **** = p < 0.0001.

After being endocytosed into neighboring cells, Tau can damage endo-membranes and thereby escape endolysosomal vesicles [7,8] (Figure 2E). This step is critical as the initial entry of seeds into the cytosol is the limiting step during intercellular Tau propagation, which initiates the subsequent seeded amplification of misfolded Tau in the receiving cell. To visualize endolysosomal damage, we employed the galectin puncta assay [24]. Human galectin-3 fused to superfolder GFP (sfGFP::LGALS3) is diffusely localized in the cytosol of hypodermal cells under steady state conditions, but redistributes to damaged vesicles and forms visible puncta [24] (Figure 2E). Formation of sfGFP::LGALS3 (Gal3) puncta indicates that the spreading and accumulation of F3^ΔK281^::mCh in the hypodermis causes endolysosomal damage (Figure 2F, G). At day five of age, animals expressing F3^ΔK281^::mCh exhibited significantly greater endolysosomal damage compared to the control (Figure 2G).

To verify that the galectin puncta assay is sensitive enough to conduct a genome wide RNAi screen, we aimed to examine the knockdown of genes that have been shown to reduce endolysosomal integrity and facilitate escape of Tau [7]. However, we encountered technical problems as the pPD49.26 vector used to generate the sfGFP::LGALS3 transgenic animals had considerable overlap with the L4440 vector used to create the Ahringer RNAi library [24–26]. The backbones of both vectors are approximately 70% identical (Figure S2). As a result, knockdown of candidate genes by RNAi caused the simultaneous silencing of the sfGFP::LGALS3 reporter construct, a phenomenon previously described as RNAi-induced transcriptional gene silencing [27]. Therefore, we generated a new sfGFP::LGALS3 strain using a different vector backbone (pDestR3-R4II).

To compare the sensitivity of the new and original sfGFP::LGALS3 transgenic strains, both were treated with chloroquine, a lysomotropic agent known to trigger lysosomal rupture [28]. Both strains showed robust, dose-dependent Gal3 puncta formation upon treatment (Figure S3). Notably, the newly generated strain displayed significantly more chloroquine-induced puncta, indicating enhanced sensitivity in detecting hypodermal endolysosomal vesicle rupture.

The newly generated strain was subsequently crossed with F3^ΔK281^::mCh and mCh animals to finally investigate the effects of ESCRT-III complex components *vps-20* and *did-2*, which are known to function in endolysosomal membrane repair and play a role in endolysosomal escape of Tau seeds [7] on Gal3 foci formation. As expected, knockdown of these genes led to a significant increase in Gal3 foci formation, indicating endolysosomal rupture (Figure 2H, I). While expression of F3^ΔK281^::mCh alone only marginally induced endolysosomal rupture, it notably enhanced the overall number of hypodermal Gal3 puncta after knockdown of the ESCRT-III components compared to the mCh control (Figure 2H, I). Thus, the chronic transmission of F3^ΔK281^::mCh appears to sensitize the hypodermal endolysosomal system to additional insults.

### Genome-wide screen for genetic modifiers of Tau-induced endolysosomal damage

Damage to endolysosomal vesicles enables the escape of transmitted amyloidogenic seeds to the cytosol, where they can promote the aggregation of native Tau proteins [10,11], potentially exacerbating disease progression. To enhance our understanding of the cellular pathways that counteract this crucial step in the propagation of Tau misfolding, we sought to identify genes that are important for the maintenance of endolysosomal integrity. Using the Ahringer RNAi library [26], 20256 bacterial clones were screened for their effect on Gal3 foci formation in the hypodermis in the presence of F3^ΔK281^::mCh in touch receptor neurons in a 96 well plate format (Figure 3A). The primary screen yielded 268 hits, which were defined as a RNAi clone that elicited at least three foci in at least two animals per well (Figure 3A). Primary hits were then validated on RNAi plates discarding all candidates that were highly toxic or did not result in ≥ 5% of animals exhibiting endolysosomal rupture on all three technical replicate plates. This resulted in 59 confirmed hits from the genome-wide screen (Figure 3B).

**Figure 3.**
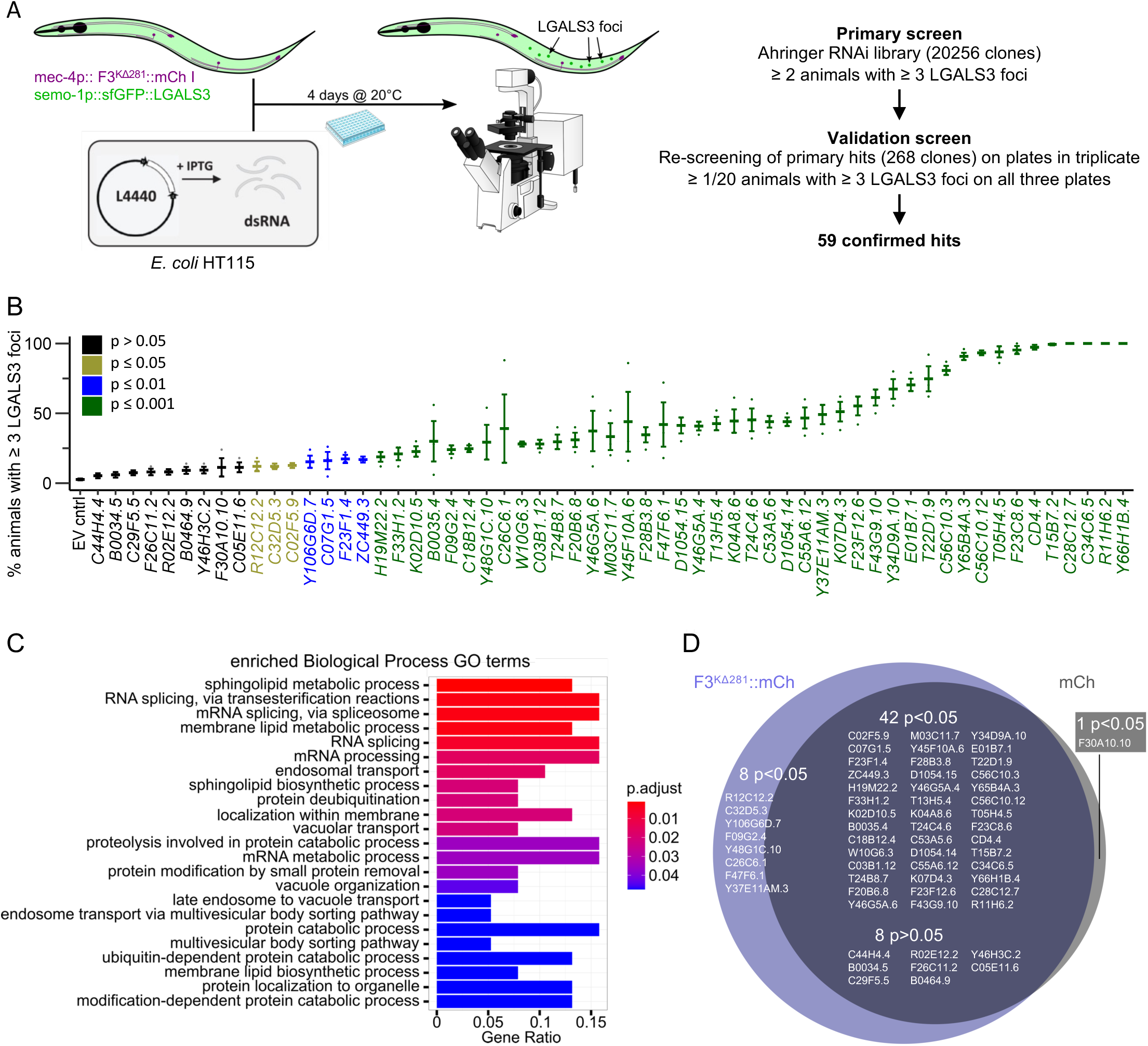
Genome-wide RNAi screen for enhancers of endolysosomal rupture in *C. elegans*. (A) Schematic illustration of the screening set-up. Age-synchronized animals expressing F3^ΔK281^::mCh in touch receptor neurons and the sfGFP::LGALS3 reporter in the hypodermis were fed HTT115 E.coli clones containing the L4440 plasmid. Upon IPTG induction, double-stranded RNA complementary to a specific *C. elegans* gene was produced, leading to targeted gene knockdown. After growth for four days at 20°C, animals were screened for Gal3 foci. The primary screen was conducted in a liquid culture in 96-Well plates. 20256 individual bacterial clones were screened. RNAi clones were considered preliminary hits if two or more animals out of 15 exhibited three or more Gal3 foci. A total of 268 preliminary hits underwent rescreening on RNAi plates. Hits were confirmed if all three plates contained at least one animal with three or more Gal3 foci after four days of growth at 20°C, resulting in a final hit list with 59 genes. (B) Quantification of animals with three or more Gal3 foci. Data are represented as mean % ± SEM. Each point represents a technical replicate. N = 3 with 10-50 animals scored per plate and replicate. Statistical analyses were conducted using Two-Way mixed-model ANOVA on rank-transformed data, with pairwise comparisons of estimated marginal means with FDR correction for multiple comparisons. Color-coding corresponds to p-values as indicated in the figure legend. (C) Enriched biological process GO terms of the significant hits in (B) using the clusterProfiler package of Bioconducter in R. (D) Venn diagram illustrating the overlap of genes identified as significantly different (42) or not significantly different (8) compared to the EV control in both F3^ΔK281^::mCh and mCh expressing animals. Eight genes were uniquely identified as significantly different in F3^ΔK281^::mCh expressing animals but not in the mCh strain, whereas one gene was exclusively detected as significantly different in mCh expressing animals, but not in the F3^ΔK281^::mCh background.

To explore the relevance of these hits, the corresponding human orthologs were identified using Ortholist2 and uniprot protein blast [29]. For the 53 of the 59 *C. elegans* genes that had at least one human homolog, GO-term enrichment analysis was performed, using the PANTHER classification system [30]. The GO-term analysis revealed an enrichment of genes belonging to specific pathways, some of which have well-established links to the endolysosomal system, such as the ESCRT complex. Among these hits were our positive controls, *vps-20* (*Y65B4A.3*) and *did-2* (*F23C8.6*), confirming the robustness of the screen. However, we also identified pathways that are linked to neurodegeneration but whose causal relationship to the maintenance of endomembrane integrity is less well understood, like the ubiquitin-proteasome system, mRNA splicing and sphingolipid metabolism (Figure 3C). The high rate of conservation among the hits in conjunction with the fact that several factors are known to be associated with the endolysosomal escape of Tau, suggests that the newly identified genes may be relevant to tauopathies.

We next asked whether the endolysosomal rupture observed for the identified hits was dependent on the continuous transmission of the Tau F3 fragment from touch receptor neurons to the hypodermis, or whether these genes act independently of Tau. To test this, we rescreened the hits using the mCh control strain. Interestingly, most hits significantly increased Gal3 foci formation compared to the empty vector (EV) control even in the absence of the F3 Tau fragment (Figure S4). Of the identified genes, 42 showed significant differences from the control in both F3^ΔK281^::mCh- and mCh-expressing animals, while eight genes were not significant in either strain (Figure 3D). However, eight genes were uniquely significant in F3^ΔK281^::mCh-expressing animals, but not in the mCh strain, indicating that the continuous transmission of the F3 Tau fragment exacerbates endolysosomal damage, leading to more genes becoming significantly different compared to the EV control. Conversely, one gene was significant in mCh-expressing animals but not in the F3^ΔK281^::mCh background. However, this discrepancy is likely due to greater variability in the formation of Gal3 foci in the F3^ΔK281^::mCh strain, which caused it to just miss the significance threshold.

In conclusion, this experiment demonstrated that while the chronic transfer of the F3 Tau fragment compromised endolysosomal integrity, thereby amplifying the effects of certain hits, most genes directly affect endolysosomal vesicles rather than acting indirectly through effects on F3^ΔK281^::mCh aggregation or transmission. In sum, this screen provided an unbiased examination of genetic modifiers of endolysosomal integrity.

### Tau aggregation exacerbated by knockdown of hits in hiPSC-derived neurons

Genes affecting endolysosomal integrity should facilitate seed entry into the cytosol and thereby promote seeded Tau aggregation (Figure 4A). Our *C. elegans* model does not express soluble Tau in hypodermal cells, which prevented us from investigating this step of misfolded Tau propagation in the nematode. To investigate whether the hits obtained from the genome-wide *C. elegans* screen affect seeded Tau aggregation, selected human orthologs were characterized in a hiPSC-derived neuronal model. These hiPSC-derived cortical neurons contain 3 mutations in the endogenous Tau locus, the E10+14 C>T and E10+16 C>T splice mutations to increase 4R Tau expression and the frontotemporal dementia-associated P301S mutation [31]. We selected 40 human genes to be tested based on their conservation and relevance in these cells, corresponding to 25 initial hit genes in *C. elegans* (Table S1). After knockdown of selected genes by siRNA, Tau-P301S/E10+14/E10+16 neurons were seeded with recombinant 2N4R P301L Tau fibrils (Figure 4B). To ensure consistent knockdown over the time course of the experiment, a second siRNA treatment was performed at mid-point (Figure 4B). The aggregation of endogenous Tau was quantified four weeks post-seeding by immunofluorescence staining [31]. The addition of fibrillar 2N4R Tau leads to the seeded aggregation of Tau-P301S/E10+14/E10+16 (Figure 4C, D). To control for different cell densities in single wells, Tau aggregation levels were normalized to the MAP2 area in the same well. Knockdown of Tau by siRNA (MAPT) served as baseline for the quantification (Figure 4C, D and S5), and wells transfected with non-targeting siRNA (nt-cntrl) were used to normalize the data obtained from three independent biological replicates (Figure S5). For all genes, knockdown efficiency was at least 50% or better, as assessed by qPCR (Figure S6). The knockdown of 8 out of the 40 genes tested (CHMP1B, CHMP4B, CHMP6, HGS, KPNB1, PSMD14, USP8, VPS37A) substantially reduced the neurite area, as seen by a more than 25% reduction of the MAP2 staining area for at least two of three biological replicates (Figure S7), thus precluding an analysis for Tau aggregation due to toxicity. For the remaining 32 genes, the knockdown of 10 genes significantly increased Tau aggregation in a meta-analysis across all three biological replicates, with additional 9 genes showing a significant increasing effect in at least one biological replicate (Figure 4E). The knockdown of one gene (SERINC4) significantly decreased Tau aggregation in the meta-analysis across all replicates, with additional two genes showing a decreasing effect in one replicate (Figure 4C-E).

**Figure 4.**
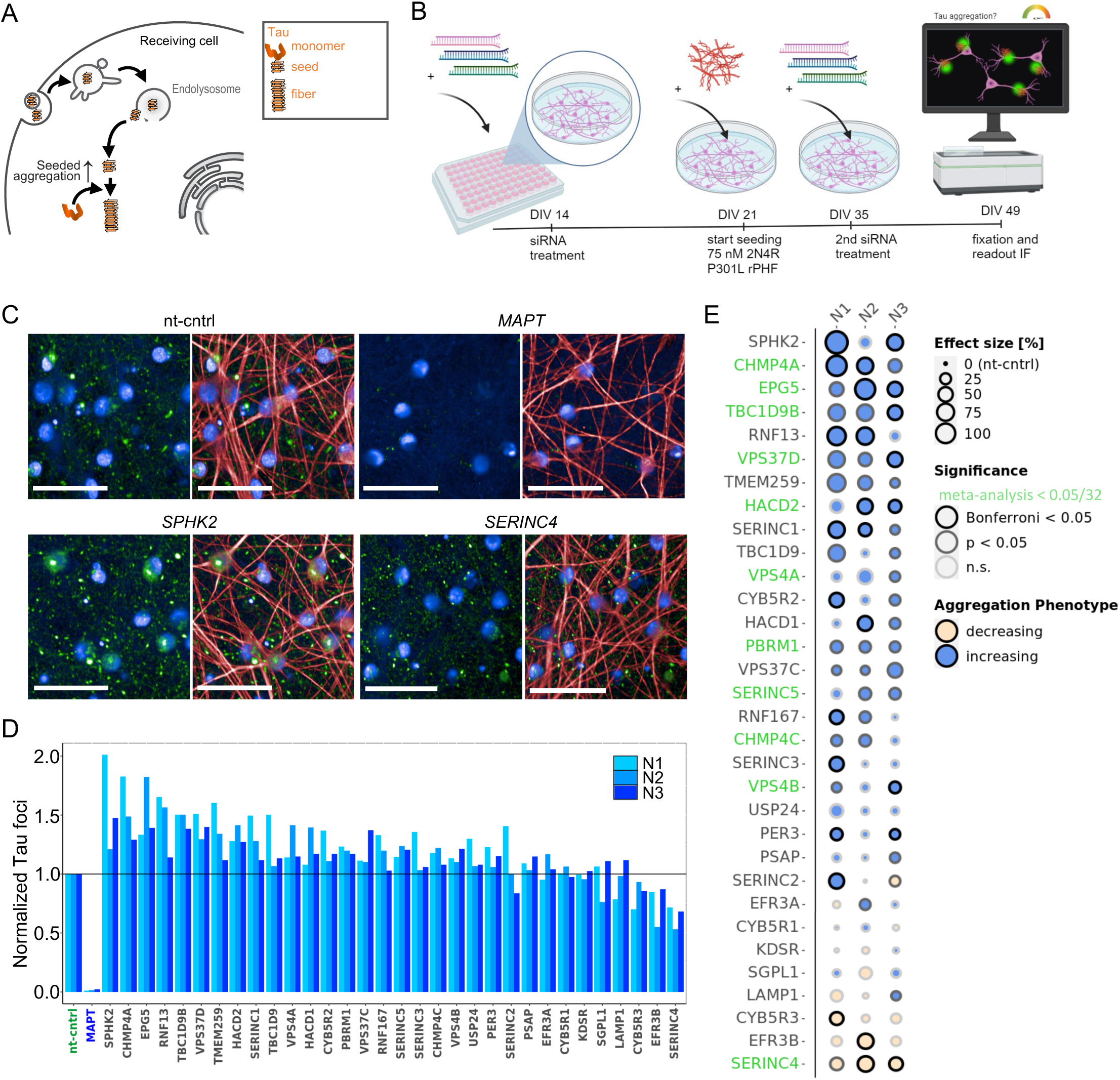
Knockdown of human orthologs of *C. elegans* screening hits modulates seeded Tau aggregation in human iPSC-derived neurons. (A) Schematic illustration of seeded aggregation of endogenous Tau following the escape of Tau aggregates from endolysosomal vesicles. (B) siRNA Tau aggregation screen timeline. Days *in vitro* (DIV) refers to the maturation age of hiPSC-derived neurons after re-plating into the final assay plate. Figure created with BioRender.com (C) Representative images of iPSC-derived neurons treated with recombinant Tau seeds and subjected to genetic knockdown using non-targeting control (nt-cntrl), MAPT, SPHK2 or SERINC4 siRNA, respectively. Scale bar: 50 µm. (D) Area of MAPT aggregate foci after knockdown of 32 non-toxic human orthologs of *C. elegans* screening hits, normalized to the non-targeting control. N1, N2, N3: Biological replicates. (E) Dot plot summarizing the effect sizes, decreasing (yellow) or increasing (blue) effects on the Tau aggregation phenotype and significance levels for each single biological replicate as well as for the overall meta-analysis per gene. Genes highlighted in green are significant in the meta-analysis after correction for multiple testing, taking all three biological replicates into account. See Table S4 for exact p-values.

We furthermore confirmed that none of the aggregation enhancers significantly altered overall Tau levels in hiPSC-derived neurons, excluding the possibility that the genes we selected may be general modulators of Tau protein synthesis or degradation (Figure S8). We also assessed whether these candidates influenced Tau aggregation in the absence of exogenous seeds. Our results showed that the hits did not significantly affect Tau aggregation without the addition of seeds, except for VPS4A, which increased unseeded Tau aggregation by approximately 2-fold (Figure S9). However, in comparison to seeded Tau aggregation, which leads to a 100-fold increase in Tau aggregation, this effect was relatively marginal.

Thus, a substantial number of these genes enhanced seeded Tau aggregation when knocked down in the hiPSC-derived cortical neuron model. This is in accordance with our expectations that increased endolysosomal rupture would lead to more Tau seeds being released from endocytic vesicles and template the aggregation of endogenous Tau-P301S/E10+14/E10+16.

To verify that these genes promote seeded Tau aggregation by inducing endolysosomal rupture, we examined their impact on endolysosomal integrity in HEK293T cells stably expressing the same Gal3 reporter used in *C. elegans* (Figure 5A). We employed an inducible CRISPR–dCas9 system in combination with a lentiviral vector expressing sgRNAs targeting the genes of interest [32]. We tested the effects of SERINC4, which significantly reduced seeded Tau aggregation, and the 10 genes that significantly enhanced seeded Tau aggregation (Figure 4E), and 3 additional genes that were nearly significant in the hiPSC screen (SPHK2, SERINC1, HACD1). Furthermore, SPHK1 was included due to its expression in HEK293T cells, as it could not be tested in hiPSC cells due to its lack of expression (data not shown).

**Figure 5.**
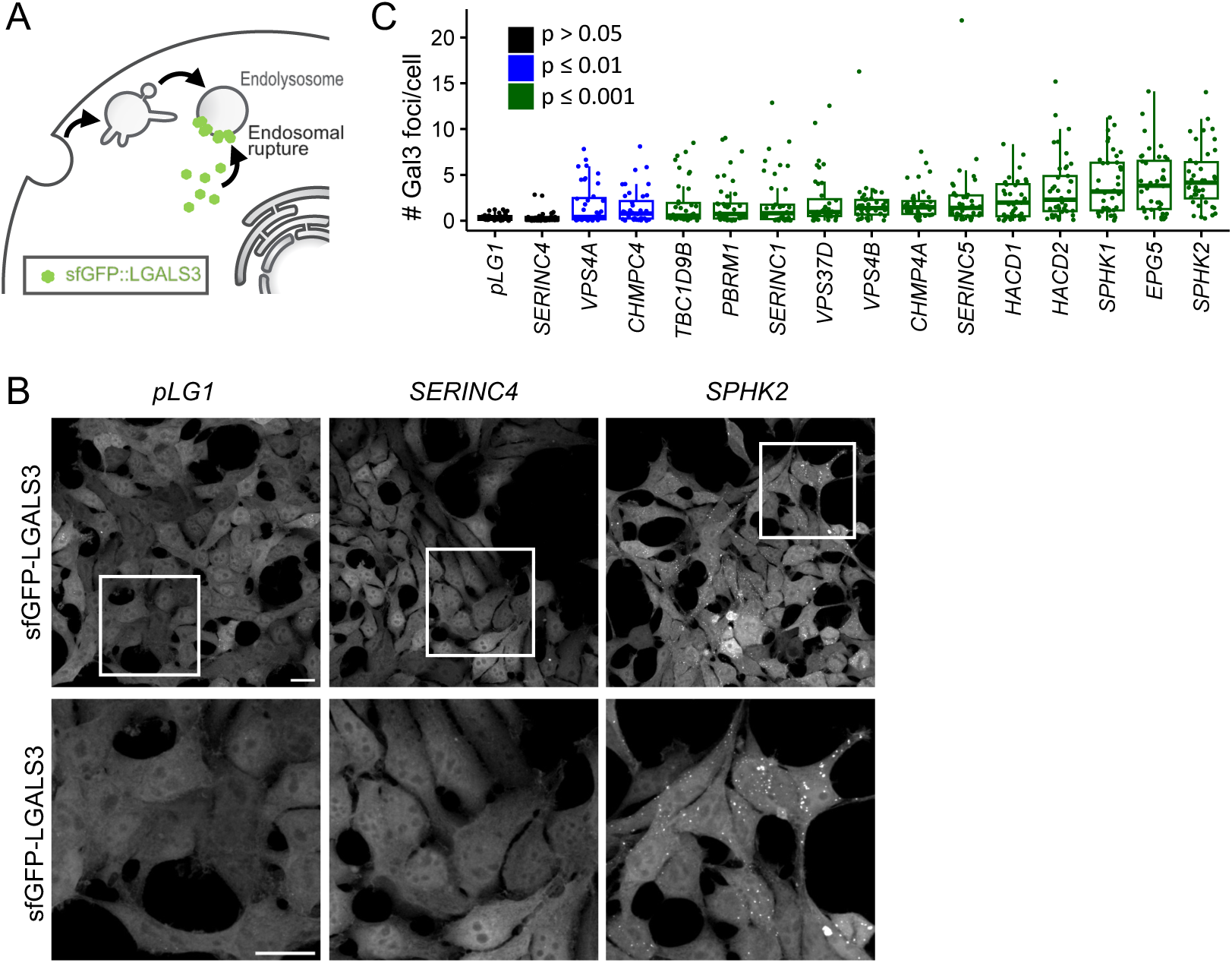
Knockdown of human orthologs of *C. elegans* screening hits modulates endolysosomal rupture in human HEK293T cells. (A) Schematic illustration of endolysosomal damage detection using the galectin-3 reporter (sfGFP::LGALS3) in HEK293T cells. (B) Representative collapsed confocal z-stacks (488 nm channel) of HEK293T cells stably expressing the sfGFP-LGALS3 reporter and the inducible dCas9 system, transiently transfected with either the empty vector control (pLG1) or pLG1 expressing sgRNAs targeting the indicated genes. Scale bar = 20 μm. (C) Quantification of Gal3 foci in HEK293T cells stably expressing the sfGFP-LGALS3 reporter and the inducible dCas9 system, transiently transfected with either the empty pLG1 vector or pLG1 expressing sgRNAs targeting the indicated genes. Downregulation of SERINC4 expression did not increase Gal3 foci formation, while targeting the expression of all other genes significantly increased Gal3 foci formation in the Gal3 reporter cells. Data are presented as boxplot, showing the mean, upper and lower quartiles, and the minimum and maximum values. Points outside the min-max range represent outliers. N = 4 with 10-11 images per gene and replicate. Each point represents one image. Statistical analysis was conducted using Two-Way mixed-model ANOVA on rank-transformed data, with pairwise comparisons of estimated marginal means with Bonferroni correction for multiple comparisons. Color-coding corresponds to p-values as indicated in the figure legend.

As expected, SERINC4 did not promote endolysosomal rupture (Figure 5B, C). In contrast, all 14 genes anticipated to enhance endolysosomal rupture significantly increased Gal3 foci formation compared to the control. These results confirm our hypothesis that these genes increase endolysosomal rupture, correlating with enhanced Tau seeding. Furthermore, these results underline the relevance of the *C. elegans* screen and the resulting hits, as the expected effects could be confirmed in two human cell models for a substantial portion of their human orthologs.

## Discussion

The gradual accumulation and progressive spreading of pathological Tau is a hallmark of tauopathies, including AD. In this study, F3^ΔK281^::mCh, a highly amyloidogenic fragment of Tau harboring a mutation linked to frontotemporal dementia was expressed in *C. elegans* touch receptor neurons. Of note, this construct serves a tool to generate a high amount of aggregated Tau. It does not fully replicate the complex pathology seen in human patients, where various isoforms of full-length and fragmented Tau species are present. Expression of F3^ΔK281^::mCh led to neuronal damage and degeneration, and a decline in mechanosensory functions (Figure 1). In addition, F3^ΔK281^::mCh was transmitted from the touch receptor neurons to the adjacent hypodermis, already starting at the first larval stage (L1) (Figure 2C, D). Although the transmission of Tau to the hypodermis gradually increased with age (Figure 2D), it occurred much earlier than in our previously described α-Syn spreading model, in which α-Syn transmission starts either on the second day of adulthood (BWM to hypodermis transmission) or the fourth larval stage (L4; DA to hypodermis transmission) [33]. This relatively early transmission of Tau compared with the α-Syn models may be due to the fact that the touch receptor neurons form neurites in the immediate vicinity of the hypodermis [34] (Figure 2B). Moreover, we utilized a truncated highly aggregation prone fragment of Tau, which seems to have a higher spreading-propensity as it is more readily found in cell culture media and the interstitial fluid of AD mouse models relative to the full-length protein [32,33].

Using this model system, an unbiased whole genome RNAi screen was performed to identify genes that impact endolysosomal membrane integrity (Figure 3A). Out of over 20,000 RNAi clones tested 59 hits were confirmed (Figure 3B), 53 of which have at least one human ortholog. This shows the power of *C. elegans* as screening model as the reduced genetic complexity allows for the identification of genes that might be masked by the redundancy of mammalian systems. Our GO-term analysis revealed an enrichment of genes with well-established links to the endolysosomal system, such as members of the ESCRT machinery [37]. Recently it was shown that the recruitment of the ESCRT-I, and -III complexes to damaged endolysosomes is essential for their repair following damage by the uptake of membrane disrupting agents [38]. As these hits are known factors linked to endolysosomal escape of Tau [7], their identification in our screen strengthens the assumption that the other newly identified genes are relevant in the context of tauopathies or other neurodegenerative diseases with a prion-like pathology. In addition to the ESCRT machinery, several hits are associated with neurodegeneration, but their specific role in maintaining endolysosomal integrity is less clear, such as the genes associated with ubiquitin-proteasome system [39].

One of the top GO-terms from the enrichment analysis was RNA splicing (Figure 3 C). Some of the most common neurodegenerative diseases, including AD, are associated with splicing defects [40]. A link between Tau pathology and splicing abnormalities has been observed in the human brain [41]. Many of the hits are associated with the spliceosomes, suggesting a general impact of splicing on Tau-mediated lysosomal rupture. However, the mechanistic link between Tau-mediated disruptions in splicing and the resulting cellular perturbations need to be further explored.

Several of the strongest hits identified in the screen were genes involved in sphingolipid metabolism. Sphingolipids are an important family of membrane lipids that are bioactive and play important roles in diverse cellular processes, including cell growth, apoptosis, and autophagy [42]. In the lipid membrane, interconnected metabolites of sphingolipids provide structural and functional properties in regard to membrane dynamics, including vesicular trafficking [43]. Our screen suggests that sphingolipid metabolism is a tightly regulated pathway, and any imbalance is problematic as knockdown of both, genes involved in *de novo* biogenesis of sphingolipids and the catabolism of sphingolipids, increases endolysosomal rupture. Disturbances in sphingolipid degradation have been identified in several genetic lysosomal storage diseases that often present clinically as progressive neurodegenerative diseases [42]. Furthermore, alterations in sphingolipid metabolism have been reported in multiple other neurodegenerative diseases, suggesting that it is an important pathway that warrants further investigation [44].

Most of the hits identified in the screen also induced significant rupture of endolysosomes in the mCh expressing control strain (Figure S4), suggesting that their effects are independent of chronic intercellular Tau transmission. Importantly, this does not diminish the relevance of these genes for AD and tauopathies. Several hits, such as those involved in sphingolipid metabolic pathways, have also been linked to the aging process. This suggests that endolysosomal leakage may be a consequence of aging [45]. Given that the escape of Tau seeds from endolysosomal vesicles is a limiting step in the spreading of Tau pathology, these genes may be relevant to disease onset and/or progression. This is supported by the fact that a considerable number of these genes also affected seeded aggregation of Tau by increasing the rupture of endolysosomal vesicles in human cells (Figure 4, 5). Moreover, the persistent transmission of the F3 Tau fragment enhanced the effects of some genes on endolysosomal vesicles in our *C. elegans* model (Figure 2H, I, 3D). Hence, chronic transmission of aggregation-prone Tau species poses a constant challenge to the endolysosomal system in the receiving tissue, which may exacerbate the damage caused by additional factors. Therefore, this strain expressing F3^ΔK281^::mCh in touch receptor neurons and the sfGFP::LGALS3 reporter in the hypodermis represents a highly sensitive model system to investigate modulators of endolysosomal rupture associated with chronic transmission of aggregated proteins.

We selected 40 human orthologs of the *C. elegans* screening hits for a siRNA screen on seeded Tau aggregation in hiPSC-derived cortical neurons. Out of the 40 genes tested, eight were toxic and thus removed from subsequent analysis. Silencing 19 of the remaining 32 human orthologs increased seeded Tau aggregation, as expected, whereas 10 genes tested had no effect and 3 had the opposite effect. One reason that some genes had no or the opposite effect could be the higher number of human orthologous counterparts of *C. elegans* genes. For example, T05H4.5 has three human orthologs, CYB5R1-3. While knocking down CYB5R2 increased seeded Tau aggregation, CYB5R1 had no impact and CYB5R3 decreased seeded Tau aggregation (Figure 5C, D). Thus, only CYB5R2 might be the functional homolog of T05H4.5 for maintaining endolysosomal integrity. Likewise, we tested five human orthologs (SERINC1-5) of R11H6.2. SERINC5 (a significant hit) and SERINC1 (with a strong tendency) both enhanced seeded Tau aggregation in hiPSC-derived cortical neurons (Figure 4), which correlates with increased Gal3 foci formation in HEK293T cells (Figure 5). In contrast, SERINC4 reduced seeded Tau aggregation in hiPSC-derived neurons without affecting Gal3 foci formation in HEK293T cells (Figure 4, 5). This suggests that the protective effect of SERINC4 is probably due to a different mechanism unrelated to endolysosomal integrity. The SERINC family is primarily involved in the incorporation of serine into membrane lipids during sphingolipid biosynthesis. However, these proteins seem to have different functions in different biological contexts [46]. For example, while all SERINC family members have been implicated in viral pathogenesis, only SERINC1 and SERINC2 are associated with Parkinsonism, and only SERINC3 is linked to melanoma [46]. The precise physiological functions of SERINC proteins remain unclear, which also complicates the interpretation of the differential effects of these proteins on Gal3 foci formation. Further studies are needed to elucidate the specific mechanisms by which SERINC family members contribute to endolysosomal integrity.

One should also consider that commercially available siRNA pools were used in this screen, and proteins with longer half-lives may only be functionally reduced at a later timepoint after the addition of recombinant Tau fibers. Weaker knockdowns of individual candidates might have resulted in an insufficient effect. Optimization of the siRNA constructs used for the knockdown could further increase the number of positive hits in these cells. Nevertheless, the silencing of a substantial number (19) of the tested genes led to an increase in seeded Tau aggregation. This effect is consistent with their ability to enhance endolysosomal rupture (Figure 5). This likely facilitates the release of greater numbers of Tau seeds from endolysosomal vesicles, which can then serve as templates for the aggregation of cytosolic Tau.

These results underscore the significance of the hits from the *C. elegans* screen, as their functional impact appears to be largely conserved in humans. This also reinforces the notion that the newly identified genes play a role in the propagation of Tau pathology and potentially also other amyloidogenic proteins in disease. Moreover, lysosomal leakage generally poses a major threat to cellular survival as it can trigger autophagy and/or lysosome dependent cell death [47]. Thus, endolysosomal integrity is not only a critical facet of prion-like propagation of misfolded proteins, but also of general cellular health. As such, further investigation of the hits identified in this study will contribute to elucidating the role of endolysosomal integrity for human health.

## Material and methods

### Maintenance of *C. elegans* and age synchronization

All animals were cultured using standard methods [48]. If not otherwise indicated, worms were grown on nematode growth medium (NGM) plates seeded with *E. coli* strain OP50 at 20°C. For RNAi, NGM medium was supplemented with Ampicillin, Tetracycline, and IPTG, seeded with the respective HT115 *E. coli* RNAi clones and grown at 20°C. Animals were age-synchronized by bleaching. Briefly, gravid adults were dissolved in 20% sodium hypochlorite solution. The surviving *C. elegans* embryos were washed with M9 buffer twice and let hatch with gentle rocking in M9 buffer at 20°C overnight. The next day, L1 larvae were distributed onto assay plates.

### Cloning of *C. elegans* expression constructs and generation of transgenic animals

The plasmid for F3^ΔK281^::mCh expression in touch receptor neurons was generated by In-Fusion cloning. Synthetic DNA coding for Tau F3^ΔK281^ and mCherry together with the *let-858* 3’UTR were ordered from Thermo Fisher scientific with *C. elegans* codon optimization using the GeneArt String software. To improve expression of the construct, three artificial introns were inserted into the mCherry ORF. To generate dsDNA fragments for directional combination, appropriate primers were designed using the Clontech In-Fusion primer design tool and employed in a PCR reaction with the respective templates. Suitable dsDNA fragments coding for the *mec-4* promoter, Tau F3^ΔK281^, mCherry and the *let-858* 3’UTR were subsequently combined with the pDest R4-R3 II vector backbone, linearized by double digestion with Sma1 and BsaA1. In-Fusion cloning according to the manufacturers protocol to create the pDest R4-R3 II_mec-4p::F3^ΔK281^::mCh::let-858 3’UTR vector. The construct for mCherry expression in touch receptor neurons was cloned by isothermal Gibson assembly. *Mec-4* promoter, mCherry tag and *let-858* 3’UTR sequences were amplified by PCR using suitable primers and combined with the pDest R4-R3 II plasmid according to the manufacturers protocol to create the pDest R4-R3 II_mec-4p::mCh::let-858 3’UTR vector. The new galectin-3 reporter construct was cloned by MultiSite Gateway cloning according to the manufacturers protocol. In brief, sfGFP::LGALS3 was amplified using suitable primers and inserted into the pDONR211 vector. Subsequently, three donor plasmids coding for the *semo-1* promoter, sfGFP::LGALS3, and unc-54 3’UTR were combined into the pDest R4-R3 II vector backbone. After each cloning step plasmids from bacterial colonies growing on selective media were extracted with the GenEluteTM Five-Minute Plasmid Miniprep Kit (Quiagen) and validated by sequencing.

Extrachromosomal arrays were created by microinjecting the expression plasmids into the gonads of young adult *C. elegans* using an InjectMan4 micromanipulator (Eppendorf, Germany). Transgenic progeny was selected using a M205 FA fluorescence stereomicroscope (Leica, Germany). To integrate extrachromosomal arrays 200 transgenic L4 larvae were irradiated in an UV Stratalinker 1800 with 90.000 µjoules. Progeny of irradiated animals were screened for 100% transmission of the transgene and outcrossed five times against the N2 Bristol wildtype strain. Primers and strains used in this study are listed in Table S2 and S3, respectively.

### Mechanosensory assay

Mechanosensory Assay was done as described in Chalfie et al. [49]. Briefly, gentle touch sensitivity was tested by stroking the tip of an eyebrow hair attached to a toothpick transversely across the posterior half of an animal. 20 worms were examined on each day for each strain in 3 biological replicates with 10 strokes each. A touch response was counted when the animal stopped or moved away from the stimulus.

### Mounting of live animals for imaging

Synchronized animals were mounted on 8-10% agarose in M9 buffer pads with a drop of mounting mix (2 % (w/v) levamisole and 50 % (v/v) nanosphere size standards (Thermo Fisher)) and covered with a coverslip.

### Scoring of PLM neurodegenerative phenotypes

Live animals were scored for neurodegenerative phenotypes of PLM neurons using an Olympus IXplore SpinSR Confocal microscope with an UplanS Apo 60x/1.30 Silicon oil objective. The neurotoxicity score for the neuronal soma was determined as follows: one or both PLM soma gone = 2; abnormal soma outgrowth = 1; no toxicity = 0. The neurotoxicity score for the PLM process was determined by adding a score of 1 for each of the following phenotypes: process branching, process break, wavy process. 10 animals were examined at each day of age in each of 3 independent biological replicates.

### Quantification of *PTL-1::mNG* and GFP fluorescence in touch receptor neurons

High resolution imaging of touch receptor neurons expressing PTL-1::mNeonGreen (PTL-1::mNG) [50] or GFP and mCh or F3^ΔK281^::mCh were obtained with a Zeiss LSM 780 confocal microscope (Zeiss, Germany) equipped with a 488 nm Argon laser (25 mW) and a 561 nm DPSS laser (20 mW), and Zeiss ZEN 2010 software. Neurons expressing GFP and mCh or F3^ΔK281^::mCh were imaged using an Olympus IXplore SpinSR Confocal microscope using an UplanS Apo 60x/1.30 Silicon oil objective. Images were obtained using a 488 nm and 561 nm laser. The corrected total cell fluorescence (CTCF) quantification was calculated from the area adjusted signal intensity minus the background signal with ImageJ software (NIH).

### Quantification of touch receptor neuron to hypodermal transmission of Tau F3^ΔK281^::mCh

The hypodermis of age-synchronized animals expressing mCh or F3^ΔK281^::mCh was imaged using an Olympus IXplore SpinSR Confocal microscope with an UplanS Apo 60x/1.30 Silicon oil objective. The 561 nm laser power was set to 30%, exposure time to 450 ms. Stacks of images with a step size of 0.35 μm were taken and the fluorescence signal (integrated density) in the hypodermis was measured using ImageJ. The fluorescence signal was normalized to the average worm size at the respective ages. The size factor was determined as normalized area of the worm in images taken with a 10x objective. 15 animals were assessed per day of age, strain and in each of 3 biological replicates.

### Quantification of lysosomal rupture in *C. elegans* (galectin puncta assay)

For quantification of animals showing hypodermal lysosomal rupture, age-synchronized animals expressing the sfGFP::LGALS3 construct in the hypodermis were seeded on OP50 or RNAi NGM plates. Animals grown in liquid culture were transferred to unseeded NGM plates for foci counting. Scoring was done using a Leica M205 FA/FCA widefield binocular microscope. 15-50 worms were analyzed per replicate.

### Chloroquine treatment

Dead OP50 (killed by multiple freeze-thaw cycles) was resuspended in S-basal (to OD_600_ 3.5) and distributed into a 96-well plate (80 μl per well). M9 supplemented with Cholesterol, Ampicillin, Tetracycline, and Fungizone containing age synchronized L1 larvae was prepared and subsequently added to the 96-well plate (15 worms per 50 μl per well). Plates were sealed with Breath-Easy foil (Diversified Biotech). In addition, plastic spacers were placed between lid and plate to enhance gaseous exchange. Two days later, Chloroquine diphosphate salt (Sigma-Aldrich) was added at indicated concentrations. Lysosomal rupture was scored in 5-days old animals. 15 animals were assessed per Chloroquine dose, strain and in each of 3 biological replicates.

### Whole genome RNAi screen on galectin puncta formation

RNAi library plates of the extended Ahringer library containing glycerol stocks of 20256 HT115 *E. coli* clones with inducible expression of dsRNA targeting the *C. elegans* genome were duplicated. HT115 *E. coli* bacteria harboring the empty vector (EV) L4440 were grown separately to be used as negative control. After overnight incubation at 37°C, bacterial dsRNA production was induced using IPTG. After 3h induction at 37°C, the plates were taken out of the incubator and cooled down to room temperature for 30 min. M9 supplemented with IPTG, Cholesterol, Ampicillin, Tetracycline, and Fungizone containing age synchronized L1 larvae was prepared and subsequently added to the RNAi plates. Plates were sealed with Breath-Easy foil (Diversified Biotech). In addition, plastic spacers were placed between lid and plate to enhance gaseous exchange. After four days of incubation at 20°C with shaking (200 rpm), animals reached the second day of adulthood and were screened for sfGFP::LGALS3 puncta. To reduce background fluorescence from remaining bacteria, the plates were placed on ice to immobilize the animals and washed once with M9. Clones were scored as preliminary hit if well had ≥ 2 animals with ≥ 3 foci. Preliminary hits were rescreened as technical replicates on three standard RNAi NGM plates and scored as confirmed hit, when each of the three plates contained ≥ 1 out of 20 animals that had ≥ 3 sfGFP::LGALS3 foci. RNAi clones of confirmed hits were subsequently sequenced to confirm/identify the target genes.

### GO term enrichment analysis

GO term enrichment analysis of hits that passed statistical significance was done in R using the open source clusterProfiler package (version 4.8.1) of Bioconductor (version 3.17) [51].

### Statistical analysis and data presentation

Statistical analysis and data presentation was done using R (version 4.4.1) with packages readxl version 1.4.3, ‘tidyverse’ version 2.0.0, ‘stats’ version 4.4.1, ‘rstatix’ version 0.7.2, ‘car’ version 3.1-2, ‘lme4’ version 1.1-35.5, ‘emmeans’ version 1.10.3, ‘dplyr’ version 2.5.0, ‘ggplot2’ version 3.5.1, ‘ggpubr’ version 0.6.0, ‘cowplot’ version 1.1.3, ‘ggbreak’ version 0.1.2 [52], ‘ggnewscale’ version 0.5.0, ‘VennDiagram’ version 1.7.3. Normal distribution of data was assessed by Shapiro Normality Test. Statistical analysis of normally distributed data was done using parametric tests, while statistical analysis of not normally distributed data was done using non-parametric tests, such as Wilcoxon signed rank test with experiments with one variable, or mixed-model Two-Way ANOVA on rank-transformed data for experiments with two variables [53]. Appropriate post-hoc tests were performed for pairwise comparisons with correction for multiple comparisons (e.g., Bonferroni or FDR adjustment). Details about the statistical tests used for each experiment and the respective results are listed in Table S4. Significance levels: non-significant (ns) = p > 0.05, ∗ = p ≤ 0.05, ∗∗ = p ≤ 0.01, ∗∗∗ = p ≤ 0.001, and ∗∗∗∗ = p ≤ 0.0001. Data presentation, sample size (number of biological or technical replicates and sample size per replicate and condition), and the applied statistical tests are indicated in the figure legends for each experiment.

### Sequence alignment and visualization

Sequence alignment was performed using msa, an R package for multiple sequence alignment (R version 3.5) [54–56]. For visualization of shared regions between aligned vector sequences, the AlignStatPlot online tool accessible at https://bioinformatics.um6p.ma/AlignStatPlot was used [57].

### Differentiation of hiPSCs into cortical neurons

The human induced pluripotent stem cell (hiPSC) lines used in this study were derived from the line SIGi00-A-9 purchased from Sigma Aldrich, and available from EBiSC (www.ebisc.org). SIGi001-A-9 carries the following mutations: MAPT IVS10+16, P301S (biallelically). The following genetic modification was additionally introduced by a CRO on behalf of AbbVie: MAPT IVS10+14 (biallelically).

hiPSCs were differentiated using a guided protocol based on a published protocol [31] with modifications. Briefly, cultures were cultured in E8 Flex (ThermoFisher) on Matrigel (Corning)-coated cell culture ware. For differentiation, cells were harvested as single cells using Accutase (ThermoFisher) and seeded at 500,000 cells per cm^2^ onto 77.5 µg/mL Matrigel-coated cell culture plasticware in E8 Flex medium supplemented with the 10 µM ROCK inhibitor Y-27632 (Millipore).

To initiate differentiation, the cells were switched the next day to E6 medium (ThermoFisher) supplemented with 500 nM LDN193189 (Stemgent), 10 µM SB431542 (R&D Systems), and 500 nM CHIR99021 (R&D Systems) for two days. On day 2, CHIR99021 was omitted and 5 µM Cyclopamine was added to the medium besides the other small molecules. Neural induction was performed for a total of 9 days and the basal medium gradually switched to N2B27 medium (50/50 Advanced DMEM-F12 / Neurobasal, 0.5% N2 supplement, 1% B27 supplement without vitamin A, 1% Glutamax, 1% Pen/Strep – all ThermoFisher) with the same supplements.

On day 9, cells were replated on 77.5 µg/mL Matrigel-coated 6-well plates in N2B27 medium with 10 µM Y-27632 at 1.5 million cells per cm^2^. The next day, N2B27 medium was supplemented with 250nM LDN193189 and with 10ng/mL FGF2 (R&D Systems) until day 15 of differentiation. On day 17, the neural rosette cortical NPC culture was prepared for freezing by dissociation into single cells with Accutase containing 10 µM Y-27632 and frozen in N2B27 supplemented with 10% DMSO (Applichem) and 10 µM Y-27632 until thawed for final differentiation.

For final differentiation, NPCs were thawed and plated on day −8 before replating on 77.5 µg/mL Matrigel-coated 6-well plates in N2B27 containing 10 µM Y-27632, 500 nM LDN193189 and 5 µM Cyclopamine. On day −5 before replating, 500 nM RO4929097 (VWR) was added once for three days to the medium to promote maturation. On the day of replating, the cells were dissociated to single cells with Accutase supplemented with 10 µM Y-27632 and replated in neural maintenance medium (Neurobasal Plus, supplemented with 2% B27+ supplement, 1% Pen/Strep, 1% GlutaMax (all ThermoFisher) supplemented with 10 ng/mL BDNF and 5 ng/mL GDNF (both R&D Systems), 200 µM Ascorbic Acid, 200µM dbcAMP, 2 µg/mL Laminin (all Sigma)) supplemented with 10 µM Y-27632 and 500 nM RO4929097 on poly-L-ornithine (0.01%; Sigma) / Matrigel coated 96-well plates (Greiner). Y-27632 was removed after 2 days after replating, RO4929097 after 5 days after plating, using 50% medium changes with neural maintenance medium from then on every 4-5 days.

### Recombinant Tau seeds

Recombinant paired helical filaments (rPHFs) were prepared by heating 40 μM human 2N4R-P301LTau monomer in PBS supplemented with 2 mM DTT to 50°C for 15 min and then cooled down to 37°C. Heparin was added to a final concentration of 40 μM, and the mixture was shaken for 6-7 days (550 rpm) at 37°C. 1 mM DTT was added daily to reduce disulfide bonds. rPHFs were centrifuged for 3 h at maximum speed in a tabletop centrifuge at 4°C to pellet the rPHFs. The supernatant (containing buffer with DTT) was removed, and the pellet was resolved in an equal volume of PBS without DTT to remove DTT and any remaining aggregation-incompetent Tau material from the seed preparation. The rPHFs were sonicated for 30 min (50% amplitude, short intervals of sonification on ice), and the concentration was confirmed using a BCA assay according to the manufacturer’s instructions. Sonicated rPHFs were aliquoted in low-binding Eppendorf tubes and stored at −80°C until later use.

### siRNA Tau aggregation screen in hiPSC-derived neurons

After the final replating, hiPSC-derived neurons were matured for 14 days before siRNA transfection. siRNA transfection using 25nmol siGenome SMARTpool siRNA (Horizon Discovery) diluted in neuronal maintenance medium (without Pen/Strep) with 0.2% DharmaFECT was performed according to manufacturer’s instructions (Horizon Discovery). After 2 days, medium was replaced with fresh neuronal maintenance medium. On day in vitro (DIV) 21, hiPSC-derived neurons were seeded with 75 nM recombinant Tau seeds (2N4R P301L rPHFs). Two days later, Tau seeds were diluted by performing a 66.66% media change with neuronal maintenance medium. On DIV 33, a second siRNA transfection was performed according to manufacturer’s instructions. 50% medium changes with neural maintenance medium were performed every 4-5 days. Neurons were cultured at least four weeks after Tau seeding until methanol fixation.

### Immunofluorescence

For analysis of insoluble Tau aggregates, hiPSC-derived neurons were washed with 1xPBS and fixed with 100% ice-cold methanol for fifteen minutes at −20 °C. After fixation, cells were washed with PBS and treated with blocking buffer (3% BSA in PBS, filtrated) for 60 min at room temperature. Primary antibodies were diluted in blocking buffer before added to the cells and incubated overnight at 4 °C. After three washes with PBS, secondary antibodies and Hoechst 33342 (H3570, ThermoFisher Scientific) were diluted in blocking buffer and incubated for 90min at room temperature while protected from light. Afterwards, cells were washed with PBS, protected from light and stored at 4 °C until imaging. The following primary and secondary antibodies were used: 1 µg/ml anti-Tau (MC1) antibody (Peter Davies), 4 µg/ml anti-Map2 antibody (Abcam, ab5392), 2 µg/ml goat anti-chicken IgY (H+L) secondary antibody, Alexa Fluor™ 647 (A21449, ThermoFisher Scientific), 2 µg/ml goat anti-mouse IgG (H+L) cross-adsorbed secondary antibody, Alexa Fluor™ 488 (A11001, ThermoFisher Scientific).

### Fluorescence microscopy and quantification

Tau aggregation in hiPSC-derived neurons was acquired and analyzed by performing high content immunofluorescence imaging conducted on either the Operetta or Opera Phenix instrument using Harmony® high-content analysis software (PerkinElmer). Imaging was performed using a 40x water objective and confocal settings with a 7-layer z-stack (0.8 µm increments). All images were displayed as maximum projections of the collapsed z-stack. Area of MC1-labeled Tau aggregates, number of Hoechst-labeled nuclei, and area of Map2-labeled somatodendritic region of neurons was quantified using Harmony® high-content analysis software (PerkinElmer). For Tau aggregation analysis, Tau aggregates were defined using MC1 staining and the Find Spot algorithm. Fluorescence intensity thresholds were set based on positive (non-targeting siRNA) and negative controls (non-seeding control) on the same plate. MC1+ spot area was quantified using the sum of the area defined by the Find Spot algorithm. MAP2 area was quantified using the sum of the area defined by Image Region. Number of nuclei were quantified using the Find Nuclei algorithm. Fragmented and condensed nuclei with area smaller than 40 µm^2 and with roundness smaller than 0.6 were excluded from further analysis.

### Data analysis and statistics of siRNA Tau aggregation screen in hiPSC-derived neurons

Quality control and data analysis for this screen was performed based on raw plate-results from Harmony High-Content Imaging and Analysis Software for n = 3 biological replicates with N = 5 technical replicates on 6x 96-well-plates each. All wells were extensively quality controlled in order to identify readouts with unexpectedly high our low outcome compared to the plate median. For such wells, images were manually assessed for abnormalities and removed if technical aberrations were observed (i.e. disruption of the cell layer due to pipetting). In total, we removed 112 gene knockdown wells and 12 non-targeting siRNA controls. Genes for which at least 2 biological replicates showed a reduction of >25% in the MAP2 area after removal of technical outliers were not considered for further analysis, as the knockdown of these genes was considered toxic for neurons.

The raw Map2 area readout was compared to the plate median to identify genes that showed systematic alterations in this readout, and parametric Welch’s t-tests with Bonferroni correction were performed to assess statistical significance for each biological replicate. To finally analyze the effect of gene knockdown on Tau aggregation, the somatodendritic area identified by Map2 staining was used to normalize the area of MC1-labeled Tau aggregates per well (Figure S2A), and plate normalization to the non-targeting siRNA control was performed by dividing each well-normalized readout by the mean of the non-targeting siRNA control per plate (Figure S2B). Parametric Welch’s t-tests with Bonferroni correction were performed to assess statistical significance for each biological replicate. To assess the significance level for each genetic knockdown across biological replicates, a meta-analysis accounting for random effects was calculated using the metacont function of meta package in R “meta: General Package for Meta-Analysis” for continuous outcome data [58]. All analyses were conducted with R version 4.1 if not stated otherwise.

### Tau protein level measurements by ELISA

hiPSC-derived neurons were cultured in 96-well plate format and transfected with siRNA as described above. After 4 weeks, cells were washed three times with 1x PBS and lysed in 50 μl/well Triton buffer (150 mM NaCl; 20 mM Tris, pH 7.5; 1 mM EDTA; 1 mM EGTA; 1% Triton-X-100; 1x cOmplete Protease inhibitors; 1x PhosStop Phosphatase inhibitors (Roche)) and diluted 1000-fold in assay buffer (20 mM NaH_2_PO_4_ pH 7.4, 140 mM NaCl, 0.05% Tween 20, 0.1% BSA) to fall within the experimentally validated linear range of the assay. ELISA plates (Maxi Sorp, Thermo Scientific) were coated overnight at 4 °C with 100 μl Tau-12 capture antibody per well (2 μg/ml; Biolegend, cat. no 806502) in 20 mM NaH_2_PO_4_ pH 7.4, 140 mM NaCl, 20% glycerol. Plates were rinsed with 250 μl/well wash buffer (20 mM NaH_2_PO_4_ pH 7.4, 140 mM NaCl, 0.05% Tween 20) and blocked for 1.5 h at RT with 250 μl/well-blocking buffer (20 mM NaH_2_PO_4_ pH 7.4, 140 mM NaCl, 0.05% Tween-20, 20% glycerol, 2% BSA). After rinsing, 100 μl/well samples were added and incubated for 2 h at RT. After further rinsing, 0.1 mg/ml Biotin-HT7 (100 μl/well; Thermo Scientific) was added as detection antibody and incubated for 1 h at RT followed by addition of Pierce Streptavidin poly-HRP conjugate (Thermo Scientific, diluted 1:10 000 in assay buffer) for 1 h at RT. TMB ELISA substrate (100 μl/well; KEM EN TEC Diagnostics) was incubated for 5 min in the dark for signal detection. The reaction was stopped by adding 100 μl/well 0.18 M H_2_SO_4_, and the absorbance was read at 450 nm on an Anthos LEDetect microplate reader (Anthos Mikrosysteme GmbH, Frisoythe, Germany).

### Quantification of mRNA levels after knock down by Q-RT-PCR

hiPSC-derived neurons were plated, cultured and transfected as described above. To determine knock down efficiency of individual genes 3 days after siRNA treatment, cells were washed 3 times with 1x PBS and then processed according to the Fast advance cells-to-ct Taqman Kit (Thermo). Taqman probes for all tested genes, as well as the housekeeping genes ß-Actin and GAPDH were purchased commercially (Thermo). The Q-PCR-reaction and analysis was performed using the Quantstudio 7 RT-PCR machine (Applied Biosystems).

### Cloning of mammalian expression constructs and generation of clonal sfGFP-LGALS3 and KRAB-Sp dCas9 expressing HEK293T cells

HEK293T cells were cultured in DMEM containing high glucose, GlutaMAX supplement, pyruvate (Gibco), and 10% FBS (Gibco) at 37 °C and 5% CO_2_. Cells expressing sfGFP-LGALS3 were generated as follows. sfGFP::LGALS3 was amplified from the *C. elegans* expression plasmid pPD49.26 by PCR and cloned into the pLenti PGK Puro DEST (w529-2) vector backbone (Addgene plasmid # 19068) by MultiSite Gateway cloning according to the manufacturers protocol. pLenti PGK Puro DEST (w529-2) was a gift from Eric Campeau & Paul Kaufman [59]. After each cloning step plasmids isolated from bacterial colonies grown on selective media were extracted with the GenEluteTM Five-Minute Plasmid Miniprep Kit (Qiagen) and validated by sequencing. HEK293T sfGFP-LGALS3 were generated using a 3^rd^ generation lentiviral system. Lentivirus was produced by transiently transfecting HEK293T cells with the pLenti PGK Puro DEST, MDL-gagpol, RSV-REV and IVS-VSVG at a ratio of 5:3:1:1, respectively. Cell media was changed 24 hours after transfection. Viral supernatant was collected 72 h post-transfection. After centrifugation at 2000 x g for 5 min, the supernatant containing the virus was concentrated by adding the PEG-it Virus Precipitation Solution (Systems Biosciences) followed by overnight incubation at 4 °C. The precipitated virus was pelleted by centrifugation (1500 x g, 30 min, 4°C), resuspended in DMEM containing 25 mM HEPES buffer and stored at −80 °C. The lentivirus was added to HEK293T followed by treatment with Puromycin to select successfully transduced HEK293T cells stably expressing sfGFP-LGALS3.

For inducible gene repression, a KRAB-Sp dCas9 PiggyBac-based construct (Addgene plasmid #84241) was stably integrated into the HEK293T sfGFP-LGALS3 cells. Cells seeded at a density of 1.5 × 10^5^ per well in a 6-well plate and transfected the following day with 0.5 μg PiggyBac plasmid containing dCas9 and 0.2 μg Super PiggyBac Transposase plasmid (Systems Biosciences) using Lipofectamine 2000. After 3 days, cells were diluted to 1 cell/well in a 96-well plate. Single cell clones were then screened for expression of BFP maker from KRAB-Sp dCas9 since antibiotic selection by puromycin was not possible due to HEK293T sfGFP-LGALS3 cells already being puromycin resistant.

### Knockdown of target genes by sgRNA in HEK293T cells expressing sfGFP-LGALS3 and KRAB-Sp dCas9

For repression experiments, single guide RNA (sgRNA) sequences were designed to be used with dead Cas9 targeting systems based on predicted sequence from Horlbeck et al [60]. Complementary strands of the sgRNA target sequences with Gibson overhang sequences (Table S2) were annealed using step-wise cooling from 95 °C. The pLG1-non-targeting sgRNA (Addgene plasmid # 109002) was digested by Blp1 and BstX1 restriction enzymes. The digested pLG1 backbone and the annealed sgRNA were assembled using Gibson assembly. The resulting pLG1-sgRNA vectors were sequence verified and purified for transfection into cells. Cells were seeded at a density of 2 × 10^4^ in DMEM with 10% FBS in a 24-well plate. 1 d after seeding, cells were transfected with 1 μg total of sgRNA per well using 1 µl lipofectamine transfection reagent (Lipofectamine 2000) in 50µl OptiMEM (Gibco). Transfected cells were induced 6h after transfection by treatment with 100 μM abscisic acid (ABA, Sigma) and fresh media. ABA was spiked in every 48h. On day 5, cells were passaged onto poly-L-lysine coated slides and treated with fresh ABA. On day 8, cells were fixed in 4% PFA in PBS for 5 min.

### Quantification of lysosomal rupture (galectin puncta assay) in cells

Cells were imaged on an Olympus IXplore SpinSR Confocal microscope with an UplanS Apo 60x/1.30 Silicon oil objective. To detect the number of foci in an image, a difference of gaussian blur was applied to max projections to isolate foci from cytosolic sfGFP signal. Thresholding was applied using ‘moments’ threshold on FIJI. Number of foci was counted using then the ‘analyze particles’ function. Cell number was then counted manually to obtain number of foci/cell for each image. N=4, with 10-11 images in each replicate.

## Acknowledgements

We thank Silke Druffel-Augustin at ZMBH and Susanne Jung at AbbVie for excellent technical assistance. We are also grateful to all members of the Nussbaum lab for their helpful discussion and constructive comments on the manuscript. The PTL-1::mNG strain was generously provided by Dr. Miriam B. Goodman. The strain BIJ34 and a pPD49.26 expression plasmid coding for sfGFP::LGALS3 were a kind gift of Dr. Bin Liu and Dr. Marja Jäättelä. Some strains were provided by the Caenorhabditis Genetics Center (CGC), which is funded by NIH Office of Research Infrastructure Programs (P40 OD010440). Special thanks to the late Dr. Peter Davies of the Feinstein Institute for providing the MC1 antibody, which has been invaluable to AbbVie Tau research. The antibody was received by AbbVie through a Material Transfer Agreement. We would also like to thank Washington University in St. Louis Genome Engineering and iPSC Center for performing gene editing (funded by AbbVie). Microwell and microscope icons in Figure 3 are from TogoTV (©2016 DBCLS TogoTV, CC-BY-4.0 https://creativecommons.org/licenses/by/4.0/). Biorender.com was used to create Figure 4B.

## Disclosure statement

D.C.S., P.R., T.L., J.S.R., L.G. and D.E.E. are employees of AbbVie. A.S. and T.R.J. are former AbbVie employees. The other authors declare no competing interests.

## Availability of data and materials

All data supporting the findings of this study are available within the paper and its supplemental material. *C. elegans* strains and plasmids generated in this study are available upon request.

## Funding

This work was funded by AbbVie, the Deutsche Forschungsgemeinschaft (grant SFB1036 TP20 to C.N.K.), and Alzheimer Forschung Initiative (grant #21053 to C.N.K.). M.P. was funded by the Erasmus+ program.

## Authors’ contributions

Conceptualization, C.A.S. and C.N.-K.; Methodology, C.A.S., G.U. and C.N.-K.; Investigation, C.A.S., N.M., M.P., A.S., P.R., D.C.S, L.G., T.R.J., D.E.E.; Formal Analysis, C.A.S., N.M., J.T., T.L., J.S.R. and C.N.-K.; Writing - Original Draft, J.T., C.A.S., and C.N.-K.; Writing - Review and Editing, C.A.S., J.T., N.M., M.P., A.S., P.R., T.L., J.S.R., D.C.S, L.G., T.R.J., D.E.E., and C.N.-K.; Supervision, T.R.J., and D.E.E., and C.N.-K.; Visualization, C.A.S., T.L., J.S.R. and C.N.-K.; Funding Acquisition, C.N.-K.

## List of Abbreviation

C. elegans: Caenorhabditis elegans
hiPSC: human induced pluripotent stem cell
PD: Parkinson’s disease
AD: Alzheimer’s disease
α-Syn: SNCA/α-synuclein
BWM: body wall muscle
DA: dopaminergic
ALM: anterior lateral microtubule cell
AVM: anterior ventral microtubule cell
PLM: posterior lateral microtubule cell
PVM: posterior ventral microtubule cell
sfGFP: superfolder Green Flourescent Protein
LGALS3: galectin-3
nt-cntrl: non-targeting siRNA
rPHFs: recombinant paired helical filaments
Tau: MAPT

