## Supplemental Files for "A Novel *C. elegans* Model for Tau Spreading Reveals Genes Critical for Endolysosomal Integrity and Seeded Tau Aggregation"

#### Supplemental online material

#### Figure S1

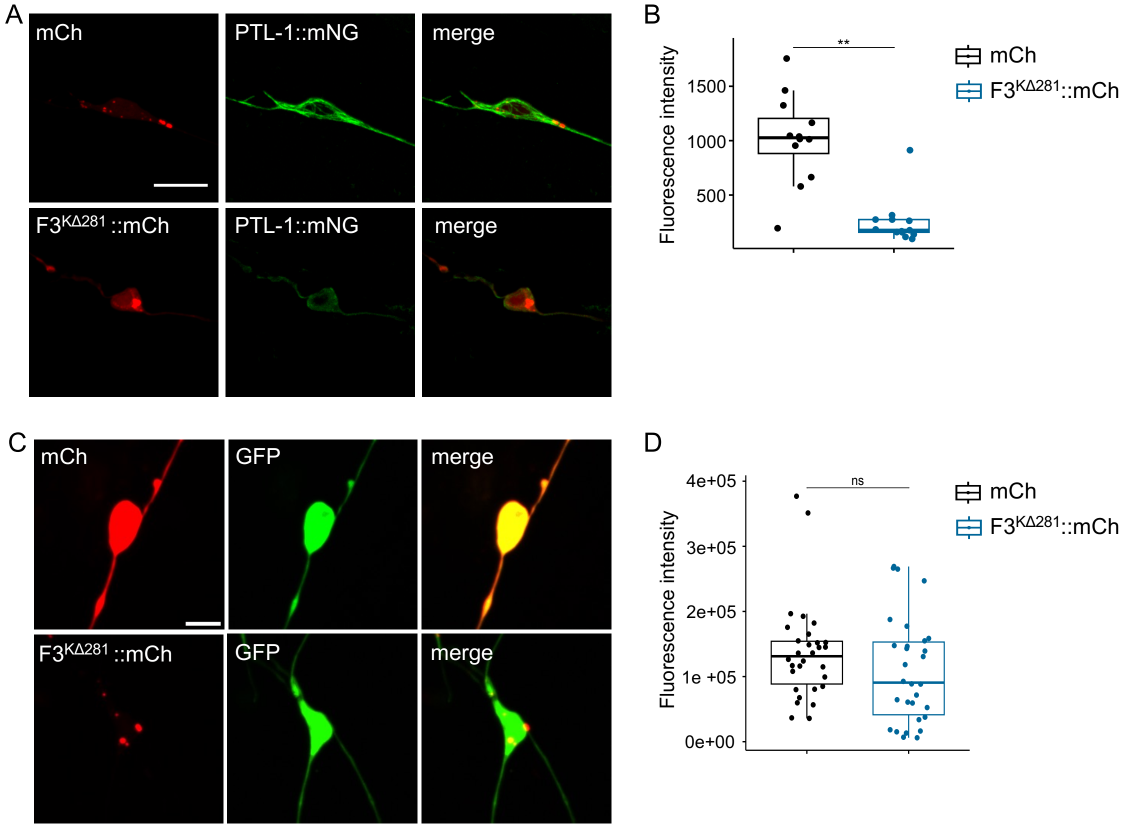


**Figure S1 Expression of F3$^{\Delta K281}$::mCh reduces PTL-1 levels in touch receptor neurons.** (A) Collapsed confocal z-stacks of touch receptor neurons of 5-days-old animals expressing PTL-1::mNG alongside either mCh or F3$^{\Delta K281}$::mCh. Expression of F3$^{\Delta K281}$::mCh in touch receptor neurons leads to reduced PTL-1::mNG signal and loss of its association to microtubules. Scale bar = 5 µm. (B) Quantification of PTL-1::mNG signal intensity in touch receptor neurons expressing mCh or F3$^{\Delta K281}$::mCh. (C) Collapsed confocal z-stacks of touch receptor neurons of 5-days-old animals co-expressing GFP alongside either mCh or F3$^{\Delta K281}$::mCh. Scale bar = 5 µm. (D) Quantification of GFP signal intensity in touch receptor neurons expressing mCh or F3$^{\Delta K281}$::mCh. The GFP signal is not affected by co-expression of F3$^{\Delta K281}$::mCh. (B, D) Data are presented as boxplot, showing the mean, upper and lower quartiles, and the minimum and maximum values. Points outside the min-max range represent outliers. Individual data points represent intensity measurements from individual animals. N = 3 with 3-10 animals per strain and replicate. Statistical analysis was done using the Wilcoxon signed-rank test. ns = not significant; ** = $p < 0.01$.

### Figure S2

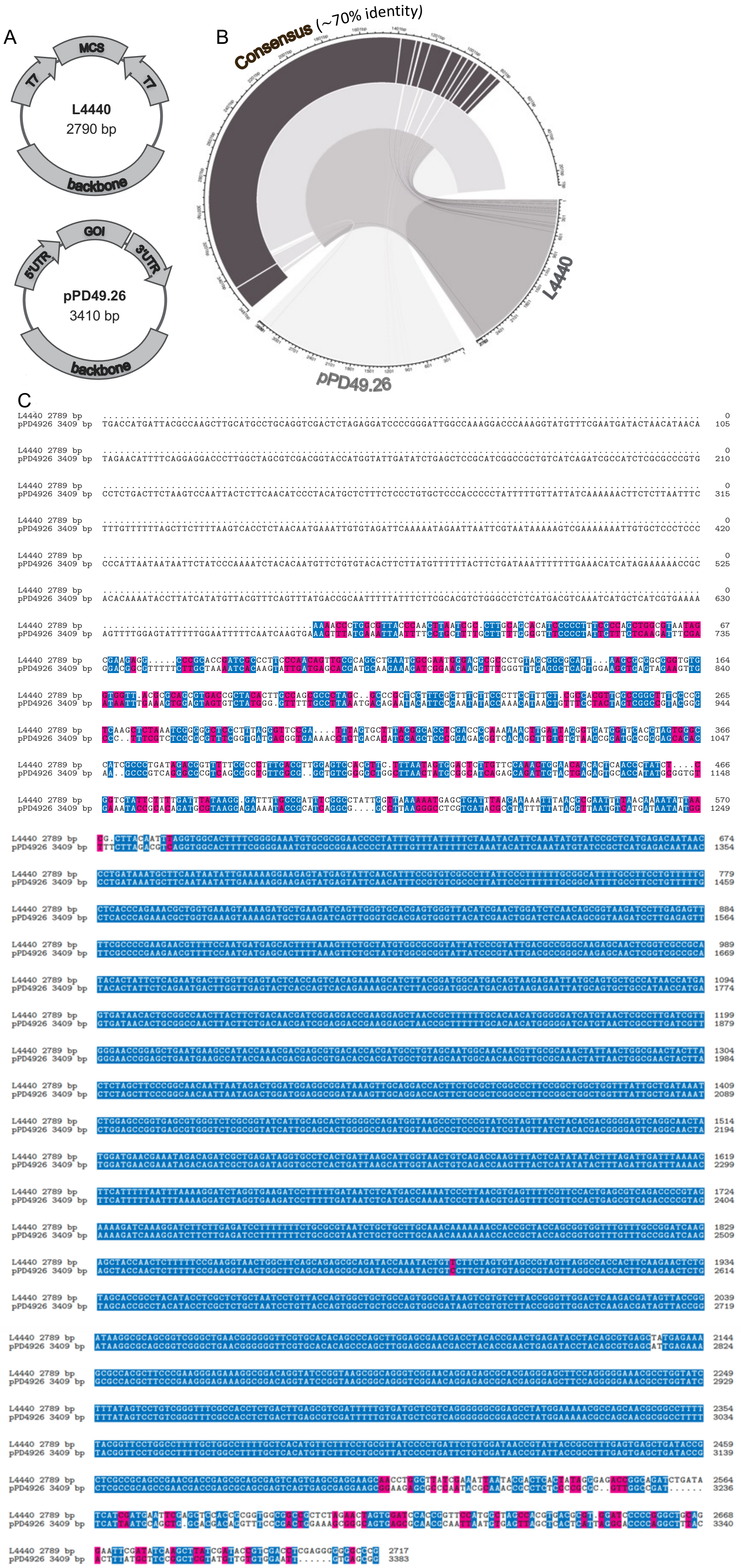
A
MCS
T7
T7
L4440
2790 bp
backbone
GOI
5'UTR
3'UTR
pPD49.26
3410 bp
backbone
B
Consensus (~70% identity)
L4440
pPD49.26
C
L4440 2789 bp
pPD4926 3409 bp

**Figure S2 The L4440 vector and pPD49.26 vector share a high degree of sequence identity.** (A) Schematic illustration of the L4440 vector and pPD49.26 vector. T7 = T7-promoter; MCS = multiple cloning site; UTR = untranslated region; GOI = gene of interest. (B) The backbones of both vectors are approximately 70% identical. (C) Sequence alignment of L4440 and pPD49.26.

**Figure S3**

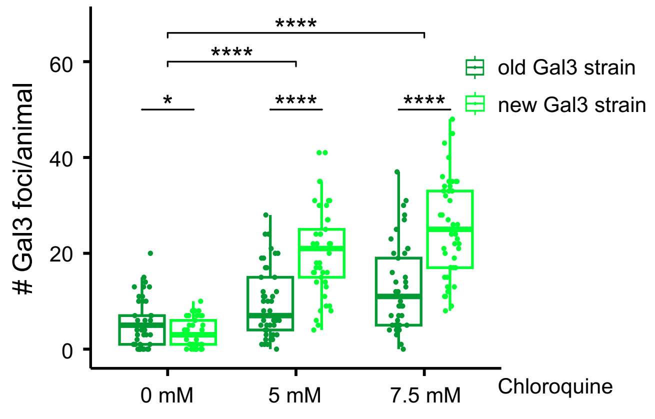


**Figure S3 The newly generated sfGFP::LGALS3 reporter strain detects endolysosomal rupture in the hypodermis with high sensitivity.** Quantification of Gal3 foci in 5-day-old animals treated with chloroquine at the indicated concentrations in both the original and newly generated Gal3 strains expressing the sfGFP::LGALS3 reporter in the hypodermis. While the original Gal3 strain shows a higher baseline of foci formation without treatment, the new strain exhibits greater sensitivity in detecting endolysosomal vesicle rupture upon exposure to the lysomotropic agent chloroquine. Each point represents the number of Gal3 foci observed in a single animal. Data are presented as boxplot, showing the mean, upper and lower quartiles, and the minimum and maximum values. Points outside the min-max range represent outliers. N = 3 with 15 animals per strain and replicate. Statistical analysis were conducted using Two-Way mixed-model ANOVA on rank-transformed data, with pairwise comparisons of estimated marginal means with Bonferroni correction for multiple comparisons. * = $p < 0.05$, **** = $p < 0.0001$.

**Figure S4**

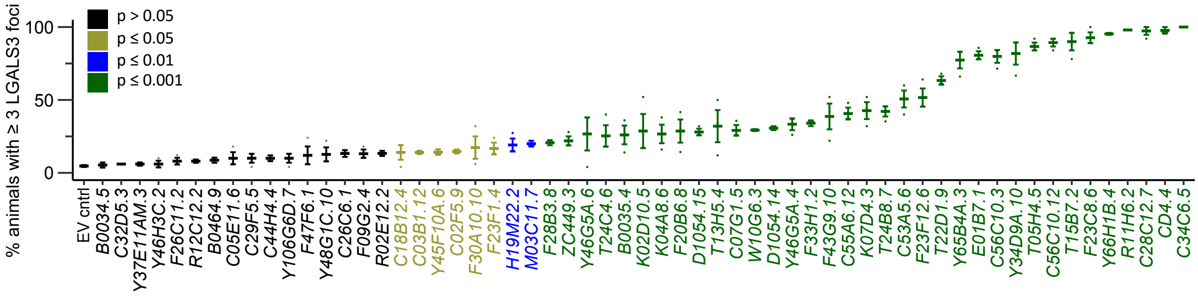


**Figure S4 Quantification of animals with three or more Gal3 foci.** Age-synchronized animals expressing mCh only in touch receptor neurons and the sfGFP::LGALS3 reporter in the hypodermis were fed HTT115 *E.coli* clones containing the L4440 plasmid for knockdown of the indicated genes. The fraction of animals with three or more Gal3 foci was determined in five-days old animals. Data are represented as mean % ± SEM. Each point represents a technical replicate. n = 3 with 10-50 animals scored per plate and replicate. Statistical analysis were conducted using Two-Way mixed-model ANOVA on rank-transformed data, with pairwise comparisons of estimated marginal means with FDR correction for multiple comparisons. Color-coding corresponds to p-values as indicated in the figure legend.

**Figure S5**

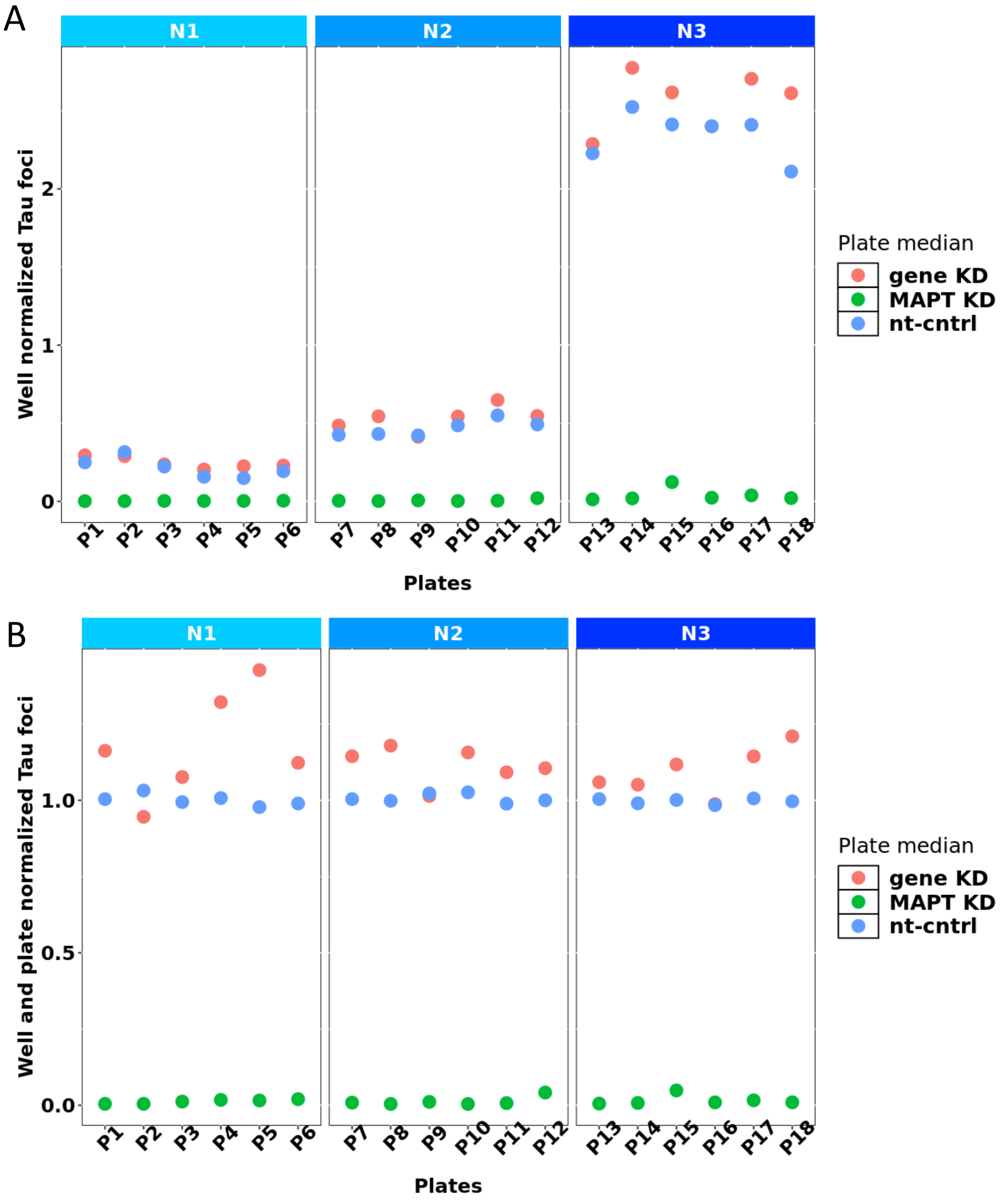


**Figure S5 Quantification data for Tau foci per plate (P1-P18) and per biological replicate (N1 – N3) in iPSC-derived neurons.** (A) Shown are the plate medians for wells with gene knockdowns (red), MAPT knockdown (green) or non-targeting control siRNA (blue). A strong difference in signal intensity was observed between biological replicates, potentially due to different batches of antibody used in the staining. (B) After plate normalization, signal intensities are comparable between biological replicates.

**Figure S6**

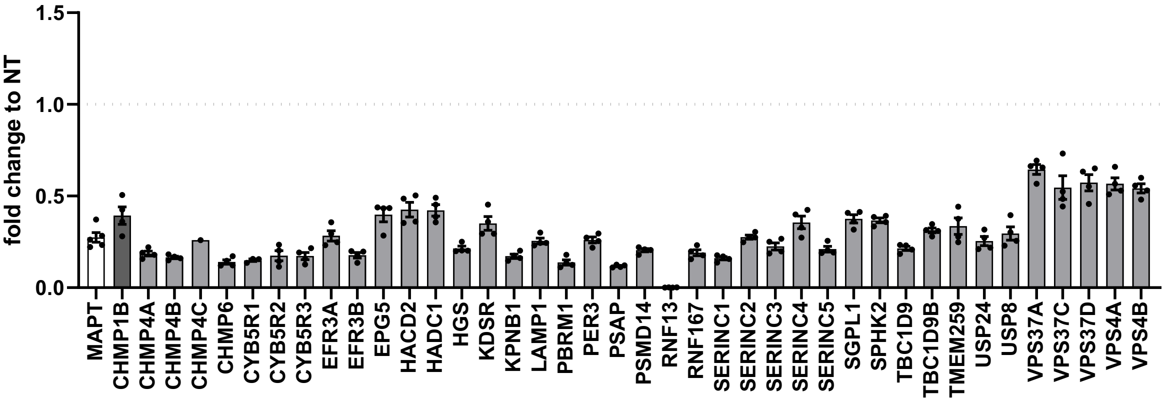


**Figure S6 Analysis of knockdown efficiency in iPSC-derived neurons.** qPCR analysis three days after siRNA treatment revealed a minimum of 50% knockdown for all genes tested. N=4 technical replicates, for CHMP4C, three replicates of siRNA-treated cells resulted in CT values >38 and could thus not be quantified. CT values for cells treated with non-targeting siRNA were all lower than 38, suggesting that despite lower expression levels, also CHMP4C siRNA treatment resulted in efficient knockdown.

**Figure S7**

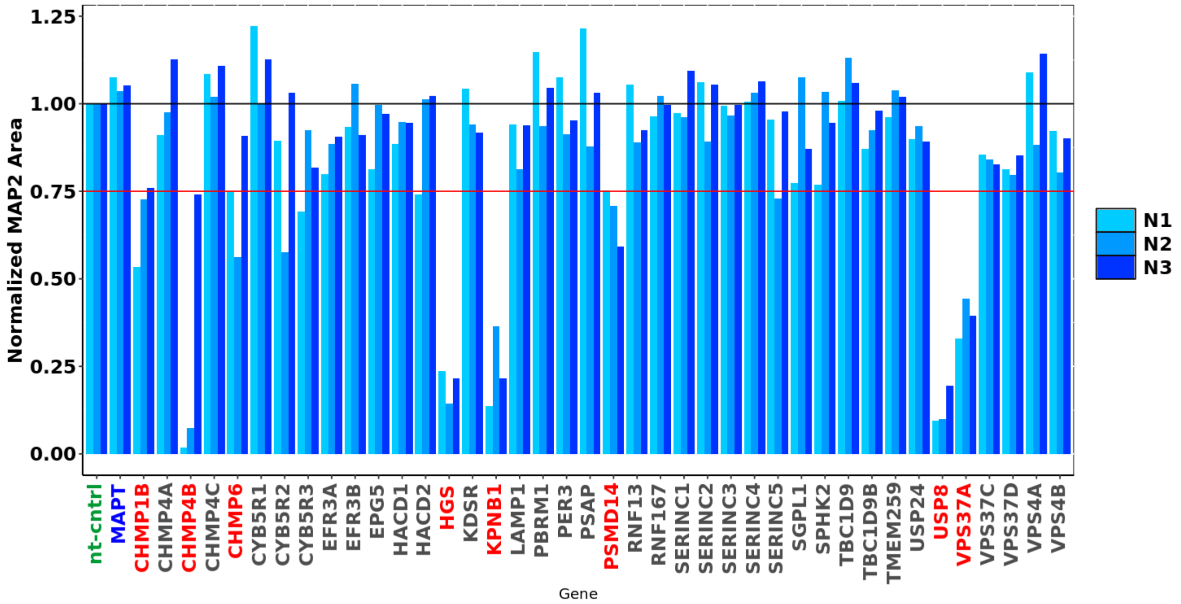


**Figure S7 Analysis of MAP2 area in iPSC-derived neurons to identify genes with toxicity after knockdown.** For each biological replicate (N1-N3), the MAP2 area was normalized to the non-targeting control. Genes showing a reduction in MAP2 area of >25% in at least 2 biological replicates were considered toxic and highlighted in red.

**Figure S8**

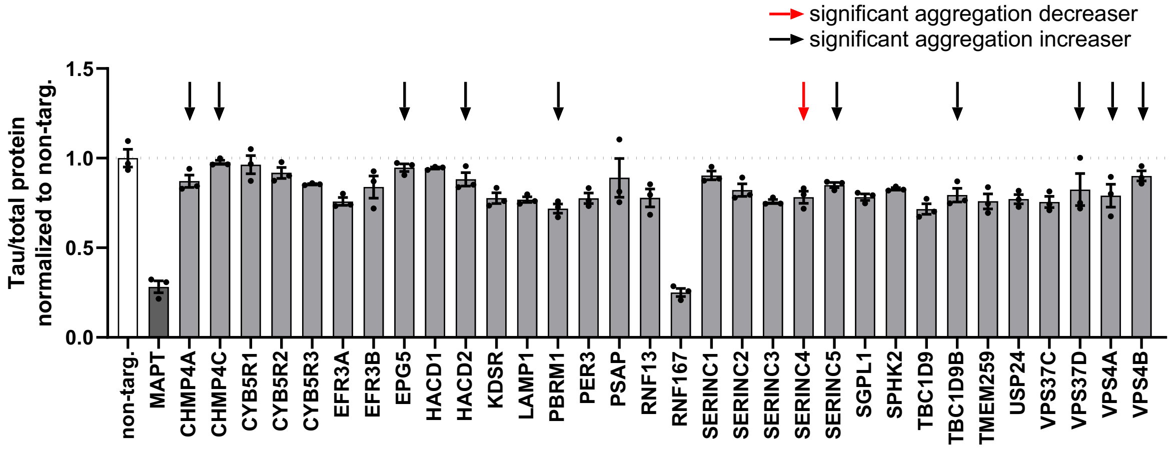


**Figure S8 Analysis of Tau protein levels after genetic knockdown in iPSC-derived neurons.** Neuronal lysates subjected to siRNA-mediated knockdown of target genes were analyzed for Tau protein levels by ELISA. While the knockdown of MAPT reduced Tau protein levels as expected, none of the genes with significant increasing effects on Tau aggregation in the meta-analysis (Figure 4E) show a corresponding increase in total Tau protein. Interestingly, the knockdown of RNF167 reduced Tau protein, however this does not lead to a reduction in Tau aggregates (Figure 4D). Additional gene knockdowns led to small, non-significant decreases in Tau levels. Data shown are representative for two biological replicates, with N=3 technical replicates each.

**Figure S9**

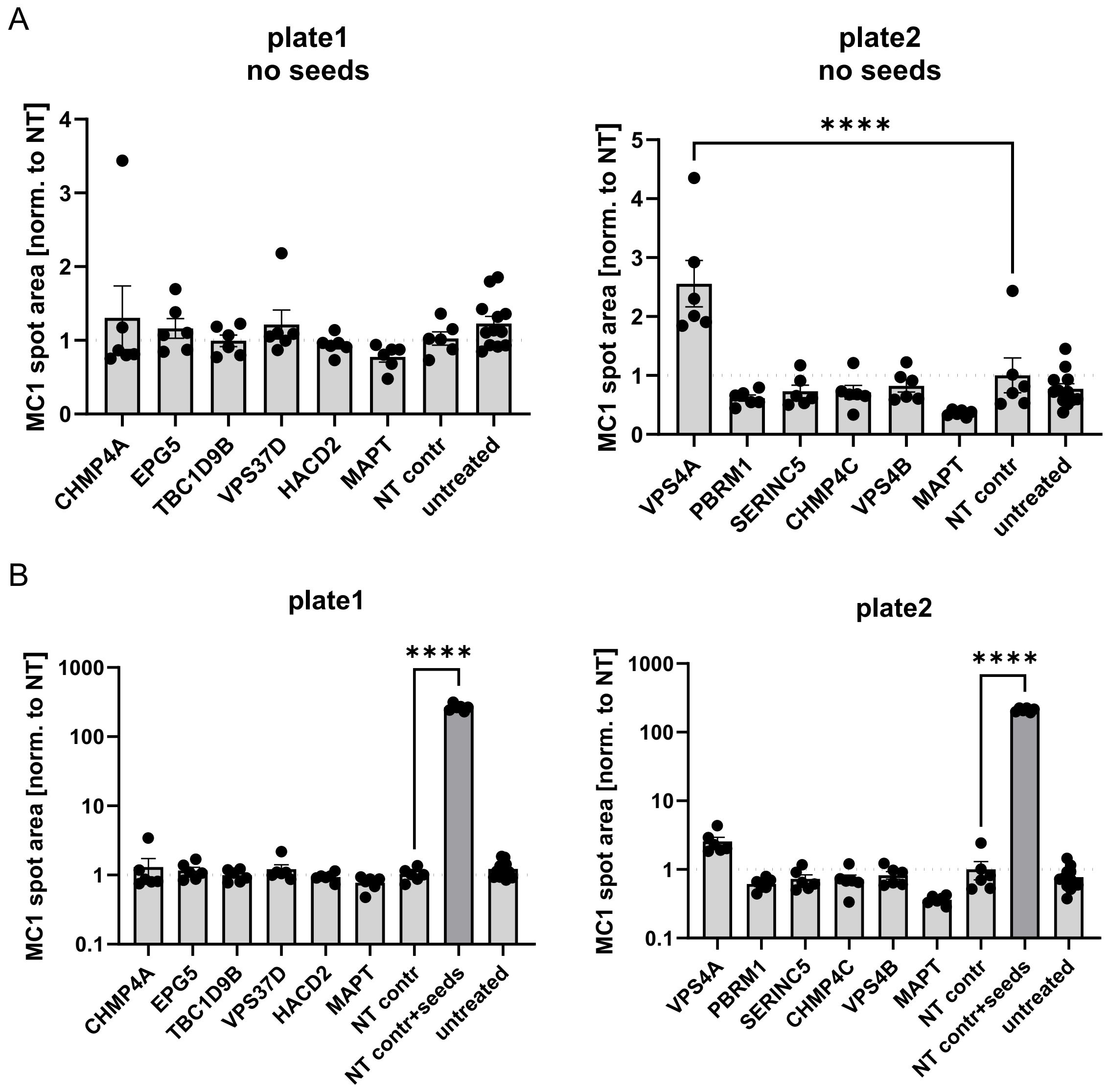


**Figure S9 Quantification of Tau foci in iPSC-derived neurons.** (A+B) Shown is the % of Tau foci for wells containing untreated cells (untreated), cells treated with non-targeting control siRNA (NT contr), or siRNA targeting the indicated genes. The knockdown of most genes did not significantly increase spontaneous aggregation of Tau. The knockdown of VPS4A resulted in approximately a two-fold increase in Tau foci formation (A); however, this effect was minimal and not significant when compared to cells seeded with recombinant Tau (NT contr+seeds), which resulted in a 200-fold increase in Tau aggregation (B).

**Graphical Abstract**

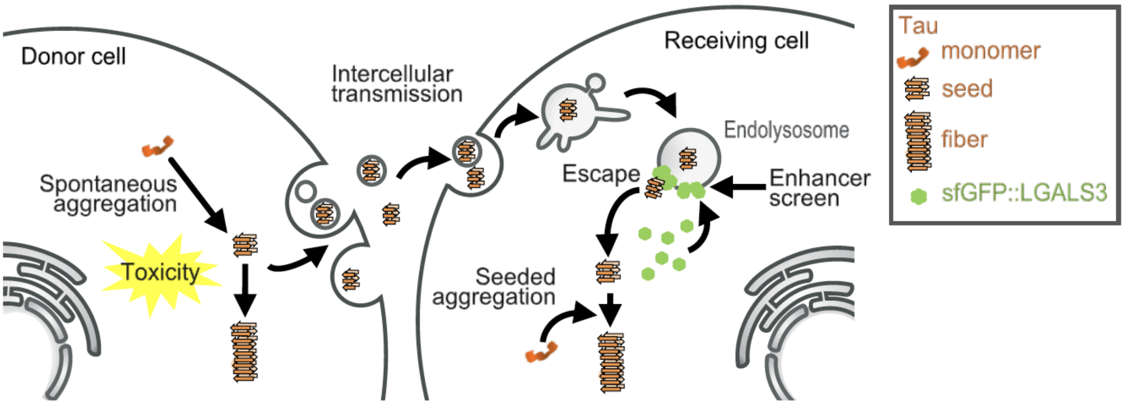


**Highlights:**

- The F3$^{\Delta K281}$ Tau fragment exhibits prion-like properties when expressed in *C. elegans*
- A genome-wide RNAi screen identified genetic modifiers of endolysosomal rupture. KD of most conserved hits increased seeded Tau aggregation in human iPSC-derived cortical neurons and induced endolysosomal rupture in HEK293T cells
- This study revealed conserved cellular pathways important for the maintenance of endolysosomal integrity: ESCRT complex, the ubiquitin-proteasome system, mRNA splicing, and sphingolipid metabolism, and shed light on the relationship between endolysosomal integrity and seeded Tau propagation
